# Multi-regional module-based signal transmission in mouse visual cortex

**DOI:** 10.1101/2020.08.30.272948

**Authors:** Xiaoxuan Jia, Joshua H. Siegle, Séverine Durand, Greggory Heller, Tamina Ramirez, Christof Koch, Shawn R. Olsen

## Abstract

The visual cortex is organized hierarchically, but the presence of extensive recurrent and parallel pathways make it challenging to decipher how signals flow between neuronal populations. Here, we tracked the flow of spiking activity recorded from six interconnected levels of the mouse visual hierarchy. By analyzing leading and lagging spike-timing relationships among pairs of simultaneously recorded neurons, we created a cellular-scale directed network graph. Using a module-detection algorithm to cluster neurons based on shared connectivity patterns, we uncovered two multi-regional communication modules distributed across the hierarchy. The direction of signal flow between and within these modules, differences in layer and area distributions, and distinct temporal dynamics suggest that one module is positioned to transmit feedforward sensory signals, whereas the other integrates inputs for recurrent processing. These results suggest that multi-regional functional modules may be a fundamental feature of organization beyond cortical areas that supports signal propagation across hierarchical recurrent networks.

## Introduction

Information processing in the neocortex involves signal representation, transformation, and transmission between processing levels or modules (Perkel and Bullock, 1968). Sensory systems are organized in anatomical hierarchies (Felleman and Van Essen, 1991; Harris et al., 2019). Areas at different hierarchical levels are thought to correspond to distinct processing modules linked by feedforward and feedback projections (DiCarlo et al., 2012; Gilbert and Li, 2013; Hubel, 1988; Kumar et al., 2010). Whereas stimulus representation (such as single neuron tuning preference and population coding) has been extensively studied (Averbeck et al., 2006; Cunningham and Yu, 2014; Hubel and Wiesel, 1962; Mountcastle et al., 1963; Orban, 2008; Parker and Newsome, 1998), the principles of spike-based transmission between neuronal populations across a processing hierarchy are much less understood (Kohn et al., 2020; Kumar et al., 2010; Zavitz and Price, 2019).

Tracking signal propagation in distributed networks requires simultaneous measurement of large numbers of interacting neurons both within and between areas in behaving animals (Buzsáki, 2004; Kohn et al., 2020; Zavitz and Price, 2019). Previous work examining inter-area communication between neurons on millisecond timescales (Chen et al., 2017; Goldey et al., 2014; Jia et al., 2013; Reid and Alonso, 1995; Semedo et al., 2019; Zandvakili and Kohn, 2015) revealed the stimulus dependence and functional specificity of pair-wise inter-regional interactions. Other studies have used local field potentials (LFPs) to measure aggregate population activity across many areas, typically focusing on gamma and beta frequency rhythms, and associate these with feedforward versus feedback signaling (Bastos et al., 2015, 2018; van Kerkoerle et al., 2014; Wong et al., 2016). However, LFP measurements cannot resolve single neurons, limiting the ability of these experiments to monitor cellular-scale signal flow. Widefield calcium imaging techniques provide simultaneous access to many cortical regions (Musall et al., 2019; Salkoff et al., 2019; Wekselblatt et al., 2016). However, these signals have much slower dynamics and don’t capture signal transmission at millisecond timescales. Thus, the larger view of network communication spanning multiple hierarchical levels of sensory processing is missing because cellular recordings with high temporal fidelity are technically challenging and have only recently become more common (Allen et al., 2019; Siegle et al., 2021; Steinmetz et al., 2021).

In our previous work, we built a recording platform using multiple Neuropixels probes to simultaneously measure hundreds of spiking neurons from up to eight levels of the mouse visual thalamo-cortical hierarchy (Siegle et al., 2021) (Fig. 1a). By analyzing correlations between spiking units we found that the average direction of the signal flow during bottom-up sensory drive follows the anatomically-defined hierarchy (Harris et al., 2019; Siegle et al., 2021). Consistent with this, we observed longer response latencies at higher hierarchy levels. However, many fundamental questions remain. Ample anatomical evidence demonstrates the existence of parallel pathways and recurrent connections between hierarchical levels of the cortex (Felleman and Van Essen, 1991; Gămănuţ et al., 2018; Harris et al., 2019; Markov et al., 2014a). In the mouse, primary visual cortex (V1) makes direct projections to all higher visual areas (Harris et al., 2019; Wang and Burkhalter, 2007), and even single cells can project in parallel to multiple areas via branching axons (Han et al., 2018). In addition, local subnetworks show clustered connectivity that could support the coactivation of groups (or assemblies) of neurons (Perin et al., 2011; Song et al., 2005; Yoshimura et al., 2005).

**Figure 1.**
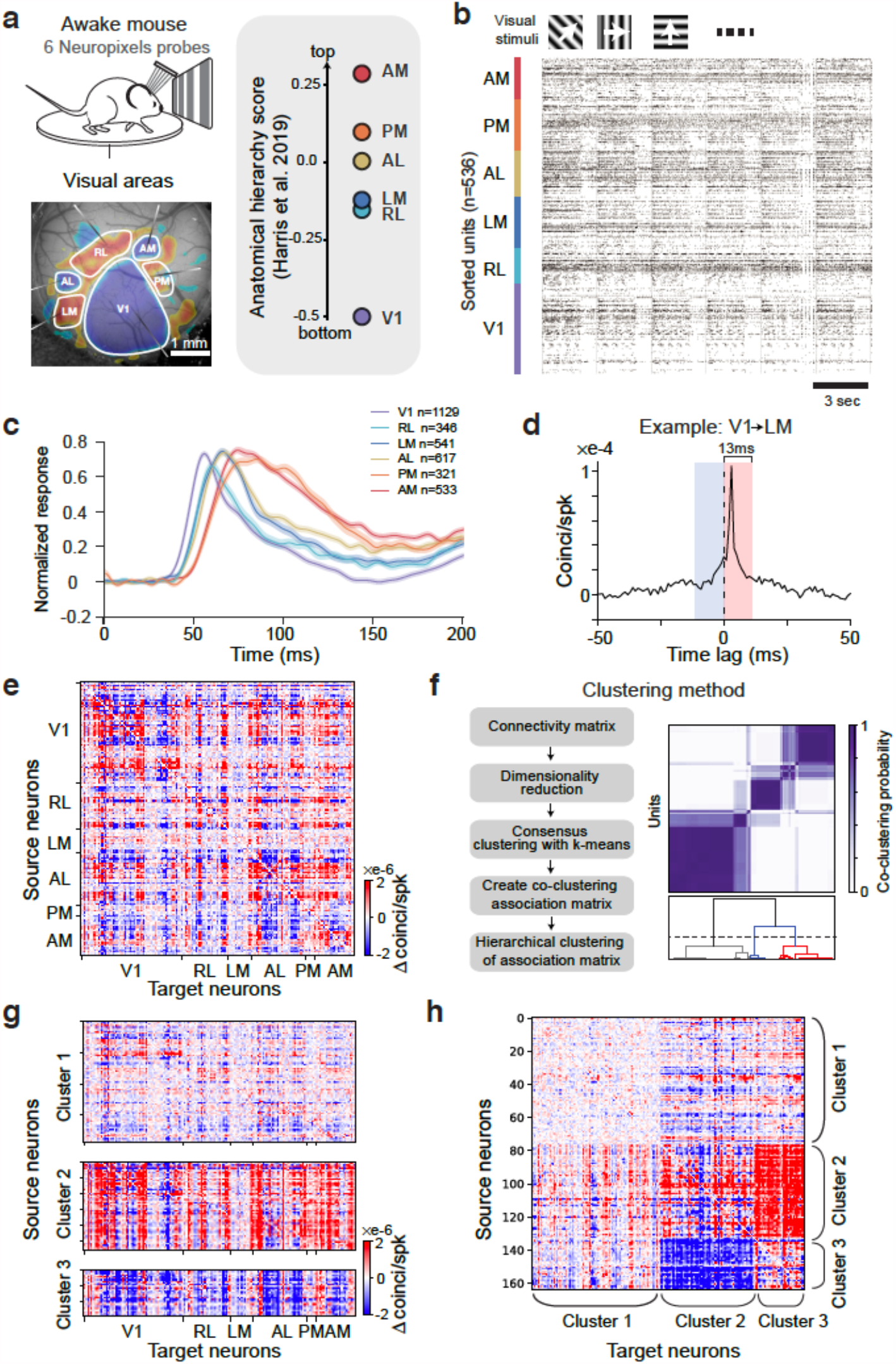
Identification of functional modules in multi-regional spike recordings. **a**) *Top*: schematic of experimental setup. *Bottom*: Neuropixels recordings from 6 visual cortical areas. Retinotopic sign map is overlaid on vasculature image to guide area targeting. *Right*: replotting of anatomical hierarchy score from *(9)* Fig. 6e. **b**) Raster plot of 536 simultaneously recorded neurons during drifting gratings stimulation (6 sequential trials are shown, each with 2 s grating presentation followed by 1 s gray period). **c**) Normalized PSTHs in response to grating stimuli from 6 visual cortical areas. **d**) Example jitter-corrected CCG between a V1 and an LM neuron with a sharp peak. Note that there is no selection criterion based on the CCG shape. A complete visualization of the raw CCGs from this example mouse is in Fig. S4. The connection weight is the difference of the CCG in a 13 ms window before and after 0 time lag, which reflects asymmetry (directionality) in the CCG. **e**) Directed cellular-scale connectivity matrix for one exemplar animal. Neurons are sorted by area and depth. **f**) Clustering procedure and result for the same data. **g**) Connectivity profiles of three functional clusters from the same mouse with source neurons organized by area and depth. **h**) Adjacency matrix organized by clusters indicating modular structure.

Given the complexity of local and inter-area connections, identifying the relevant signal transmission modules is challenging. The canonical interpretation is that each cortical area corresponds to a processing stage that performs local computations and sends output signals sequentially from one area to the next. However, this standard view is likely only a first-order characterization. Because of dense lateral and recurrent connectivity, it is possible that major communication hubs actually span multiple anatomical areas. Thus, a critical foundational step is to subdivide the system into functionally relevant processing modules at the multi-regional cellular level.

Here, we take such an approach by decomposing the cortical network into functional modules. Using multi-area recordings from six hierarchical levels, we infer signal flow based on the statistics of leading-lagging spike timing between neurons. Treating each neuron as a node, we create an adjacency matrix with directed weights and then use an unsupervised clustering algorithm to uncover multi-regional sets of neurons based on their shared functional connectivity patterns. This method revealed two major visual-engaged modules that spanned the cortical hierarchy. We find consistent evidence suggesting that one module is mainly involved in the feedforward process to distribute sensory information, while the other is more engaged in the recurrent process to integrate information.

## Results

We used up to six Neuropixels probes to record populations of neurons from six areas of the mouse visual cortex. Each area has its own map of visual space (Garrett et al., 2014), and resides at a different hierarchical level as determined both anatomically (D’Souza et al., 2020; Harris et al., 2019) and physiologically (Siegle et al., 2021). Area V1 is at the bottom of the hierarchy, followed by areas RL/LM, AL, PM, and AM (Fig. 1a). Overall, recording sessions in this study yielded 632 ± 18 simultaneously recorded neurons per experiment (a.k.a. sorted units (Siegle et al., 2021)) distributed across cortical layers and areas (*n* = 19 mice, mean ± SEM) (Fig. 1b). We used full-field drifting grating stimuli to provide a strong bottom-up sensory input to the system to evoke a large number of spikes recorded per unit time for our functional connectivity estimation. Consistent with the known visual hierarchy in the mouse (Harris et al., 2019; Siegle et al., 2021), the mean response latency in each area followed a sequential progression (Fig. 1c). However, all areas were co-active for substantial portions of the visual response, thereby providing opportunities for recurrent interactions. To facilitate functional connectivity analysis, neurons in our dataset were filtered by minimal firing rate (>2 spikes/second) and receptive field location (center of RF at least 10 degree away from the edge of the monitor). After filtering, *n* = 3487 units remained, which is 29% of total units recorded across all mice (see Methods for details).

### Identification of multi-regional functional modules

To characterize fast timescale interactions relevant to signal transmission, in each mouse we quantified spike timing cross-correlations between all pairs of neurons using jitter-corrected cross-correlogram (CCG) analysis; this captures relative spike timing between two neurons within the jitter window (25 ms) but removes stimulus-locked signals and correlations longer than the jitter window (Jia et al., 2013; Smith and Kohn, 2008a). For each neuronal pair, we determined their connection weight by computing the difference of the CCG in a 13 ms window (half of the jitter window) before and after 0 time lag (Fig. 1d), which is an indicator of the asymmetry of the CCG with short delays. Note that we didn’t select CCGs based on their shape (e.g. sharpness of the peak), so this measure is likely to include both mono- as well as polysynaptic connections. The sign of this weight describes the *signal flow direction* (temporally leading or following) between pairs of neurons. By computing the connection weight for all pairs, we produced a connectivity matrix describing the directed functional interactions of all simultaneously recorded neurons (Fig. 1e). These functional connectivity matrices displayed non-random structure (Fig. S1). Inspection indicated there existed separable groups of neurons with similar patterns of functional connectivity. We next sought to algorithmically uncover these groups by clustering their connection profiles.

To systematically identify sets of source neurons with similar connections, we clustered the interaction matrix by treating connections from each source neuron to all neurons in the recorded network as features (Fig. 1e,f; Fig. S2). This procedure yielded three robust clusters of source neurons (Fig. S3): cluster 1 had mostly weak connections; neurons in cluster 2 were dominated by strong positive connection weights indicating they tended to lead (or drive) network activity; and neurons in cluster 3 were dominated by strong negative connection weights indicating they are mainly driven by and follow activity in the network (Fig. 1g, h). These three clusters were consistently identified in each mouse we examined, suggesting they are a core organizational feature in the mouse visual cortical network (Fig. 2a). Given the bias in positive versus negative connection weights from each cluster (and its functional implication of directionality), we refer to cluster 2 source neurons as the ‘driver module’ and cluster 3 source neurons as the ‘driven module’ (Fig. 2b). Supporting the robustness of these clusters, we observed similar network modules using spectral clustering and bi-clustering algorithms (Pedregosa et al., 2011) (Fig. S3). These modules are readily apparent during periods of high-contrast visual stimulation, but are barely visible during spontaneous activity (gray screen; Fig. 2c,d).

**Figure 2.**
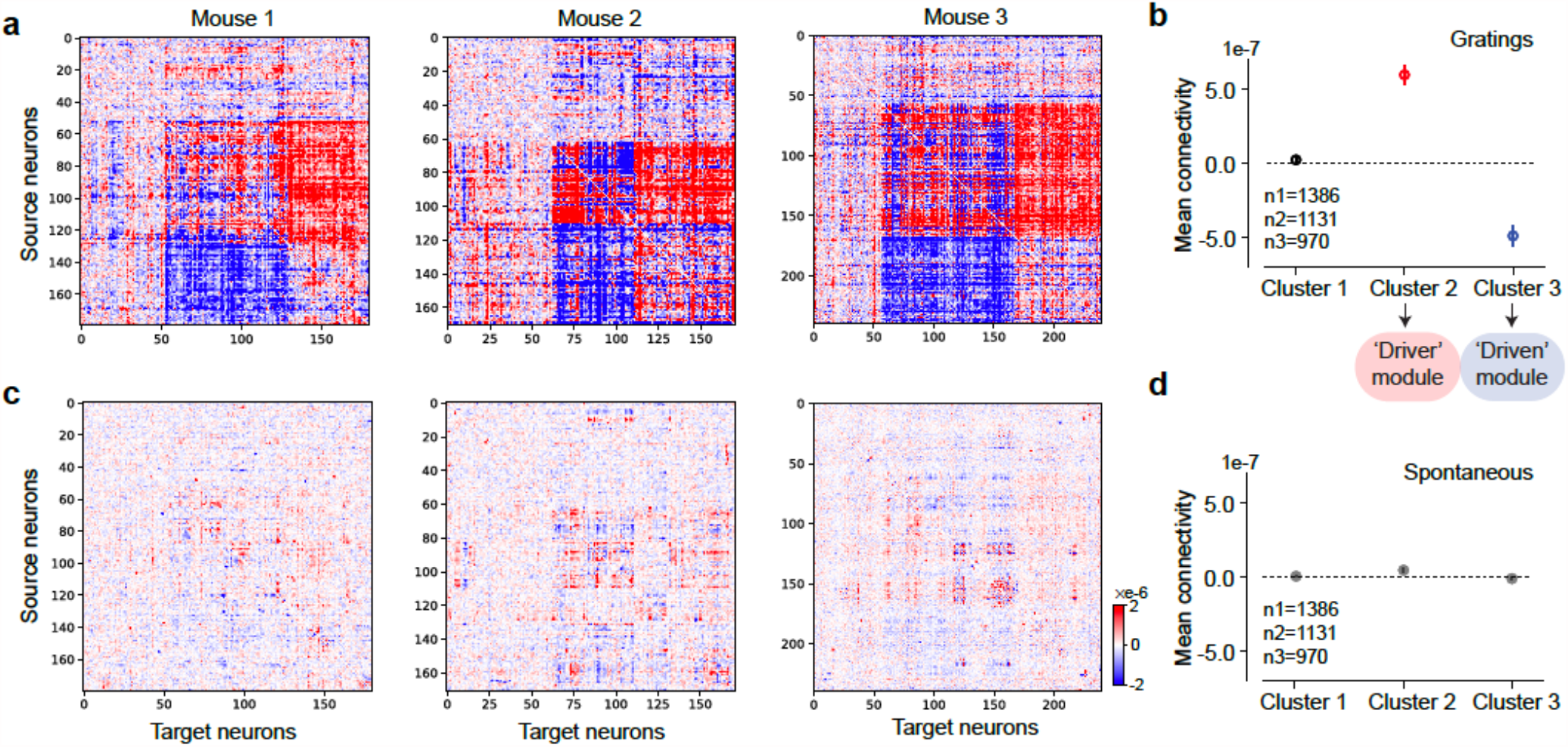
Consistent directed functional modules observed across mice. **a**) Adjacency matrices of three example mice organized by modules during drifting grating. **b**) Functional connection strength averaged over all n=3487 neurons in each cluster across all n=19 mice for drifting gratings. Mean connection strength of cluster 1 is not significantly different from 0 (Two-tailed 1-sample t-test, T=2.3, p = 0.02). Cluster 2 is significantly positive (stats=17.8, p = 1.3e-62). Cluster 3 is significantly negative (T=-14.6, p = 1.5e-43). **c**) Adjacency matrices of the same three example mice organized by modules during spontaneous activity. **d**) Functional connection strength averaged over all neurons in each cluster across all mice during spontaneous. Mean connection strength of cluster 1 is not significantly different from 0 (1-sample t-test, T = 4.5, p = 7.5e-6). Cluster 2 is significantly positive (T = 15.9, p = 1.3e-51). Cluster 3 is significantly negative (T=-2.8, p=0.0058). Error bars represent s.e.m.

In which cortical areas and layers do neurons in these two separate modules reside? Interestingly, neurons in both the driver and driven modules were spatially distributed across all levels of the cortical hierarchy rather than being localized to specific regions (Fig. 3a). Nonetheless, the proportion of neurons in the two modules showed area biases. Overall, driver neurons decreased along the hierarchy (Fig. 3a; Spearman’s correlation with each area’s hierarchical position (Harris et al., 2019): r = -0.89, p = 0.019), whereas driven neurons increased (Spearman’s correlation = 0.89, p = 0.019 for driven module). Both driver and driven neurons were present in all cortical layers but also showed laminar biases. Driver neurons were enriched in the middle and superficial layers, whereas driven neurons were more common in deeper cortical layers (Fig 3b). Anatomical tracings reveal that neurons mediating feedforward projections tend to originate in superficial layers (Felleman and Van Essen, 1991; Harris et al., 2019; Markov et al., 2014a), with the fraction of feedforward projecting neurons in superficial layers decreasing along the hierarchy (Barone et al., 2000; Markov et al., 2014b). Neurons in the driver module followed the same pattern, suggesting this module could be involved in feedforward processing, while the driven modules might be more involved in recurrent processing.

**Figure 3.**
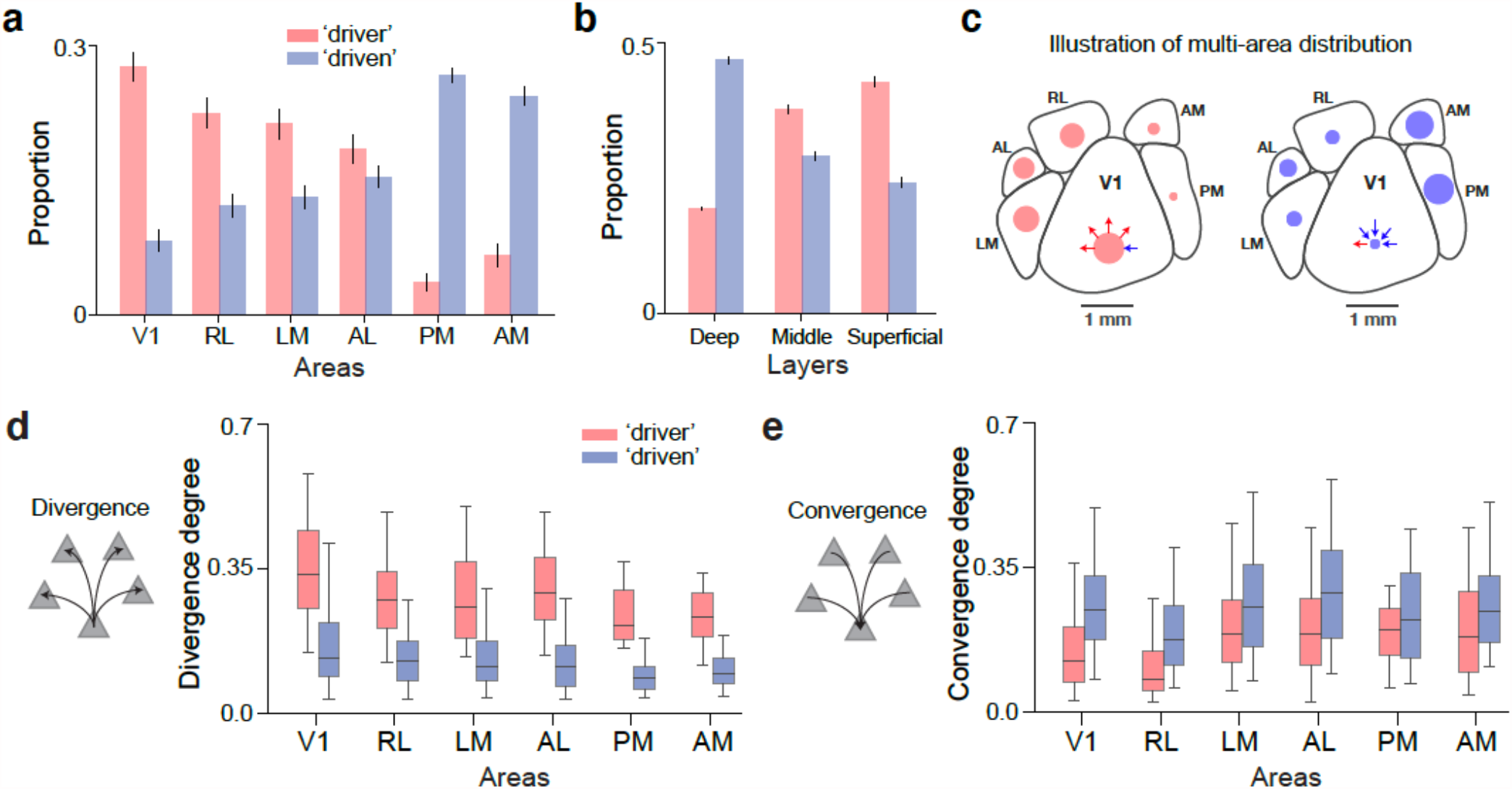
Area distribution and divergence/convergence differs across modules. **a**) Proportion of neurons in driver and driven modules across the visual cortical hierarchy. **b**) Cortical layer bias of the driver and driven neural modules for all mice. Error bars in a) and b) represent bootstrapped standard error. **c**) Illustration of the relative proportion of neurons in driver and driven modules across the visual cortical hierarchy. Red dots indicate the driver population, which has higher divergence. Blue dots indicate the driven population, which has higher convergence. The relative size of the dots corresponds to their proportion across hierarchy shown in a). **d**) Boxplot of divergence degree distribution across areas for driver and driven modules. Box indicates first quartile to third quartile, with middle line indicating median of the distribution. The whiskers indicate 5% and 95% percentiles. Divergence degree of driver module decreased along the visual hierarchy (Spearman’s r = -0.83, p = 0.04). **e**) Convergence degree distribution for driver and driven modules. Convergence degree of driven module was not correlated with hierarchy level (Spearman’s r = -0.03, p = 0.96).

### Signal integration and distribution by different modules

Node convergence (*input* to one neuron from others) and divergence (*output* from one neuron to others) are two network properties that differentially support the integration versus distribution of signals (Tononi et al., 1998). The predominately positive connection weights of the driver neurons versus the mostly negative weights of the driven neurons suggests their convergence and divergence differs. To quantify this, we computed the divergence degree of each neuron as the number of significant positive connections (outward projections from the neuron) relative to the size of the network. We define a significant connection as those with an absolute weight greater than 10^−6^ coincidences/spike, a threshold value defined by half of standard deviation of the weight distribution across all mice (see Methods and Fig. S4). Likewise, convergence degree of a neuron was computed as the fraction of significant negative connections. Driver module neurons had higher divergence than the driven neurons (Fig. 3d; 2-way ANOVA across areas, between modules F = 1058.6, p = 3e-188; among areas F = 33.2, p = 1e-32; interaction F = 3.0, p = 0.013). In contrast, the driven module neurons had higher convergence (Fig. 3e; between modules F = 239.9, p = 5.6e-51; among areas F = 20.6, p = 3.9e-20; interaction F = 3.1, p = 0.007). Thus, from a network perspective, neurons in the driver module are better positioned to distribute information, whereas neurons in the driven module are better positioned for signal integration.

The greater convergence to driven neurons indicates they combine signals during visual processing. In the visual system, the convergence of multiple simple-like neurons that are modulated by drifting grating phase can give rise to complex-like neurons with tolerance to the different grating phases. Simple- and complex-like neurons can be quantified using a modulation index (MI) that describes the degree of modulation at the preferred temporal frequency of a drifting grating stimulus (Matteucci et al., 2019), with simple-like cells having a large MI, while complex cells have a small MI (in the limit, close to zero). To test whether the driven module contains more complex-like neurons consistent with their dominant converging inputs relative to the driver module, we computed the MI for individual neurons and compared between the two modules (Fig. 4a,b). Overall, the driven population was more complex compared to the driver population (p=3.3e-24; Mann Whitney U test). Since there are more simple like neurons in V1 (reflected by the bi-model distribution in Fig. 4c), we also tested the MI between driver and driven modules if only comparing higher-visual areas. Driver neurons have significantly higher MI compared to driven even after removing V1 (p = 0.00011; Mann Whitney U test). This result is consistent with the driver neurons’ higher convergence and suggests they are functionally deeper in the visual processing pathway.

**Figure 4.**
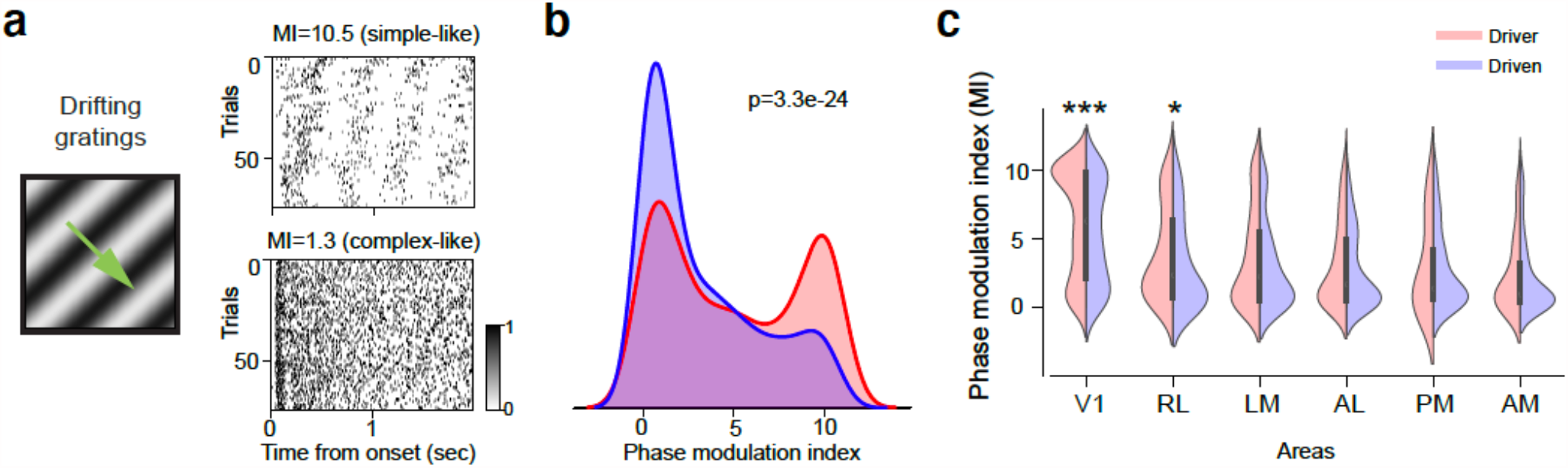
Neurons in the driven module are more complex-like compared to those in the driver module. **a**) Illustration of drifting gratings stimuli and the raster plot of two example units from driven and driven modules. **b**) Distribution of phase modulation index (MI). **c**) Mean MI of neurons belonging to different modules in different brain areas. Error bars represent s.e.m. Significant test by Mann Whitney U test. * indicates p<0.05. *** indicates p<0.001.

### Signal transmission between modules

Having now defined two robust network modules, each spanning the visual hierarchy, we next sought to evaluate signal flow within and between these separable network clusters. To do this, we investigated the weight (adjacency) matrix defined previously (Fig. 1,2), which describes directed interactions between all recorded neurons. We first focused on signal transmission between modules by examining the subnetworks defined by the connections from driver to driven neurons (Fig. 5a top) and from driven to driver neurons (Fig. 5a bottom). Connections between these modules were largely unidirectional; that is, driver neurons from each source area (including top areas of the visual hierarchy, e.g. PM and AM) made output connections to driven neurons, and driven neurons in each source area received input connections from driver neurons (Fig. 5b). The asymmetry of these connections indicates a largely unidirectional signal flow from the driver to driven module.

**Figure 5.**
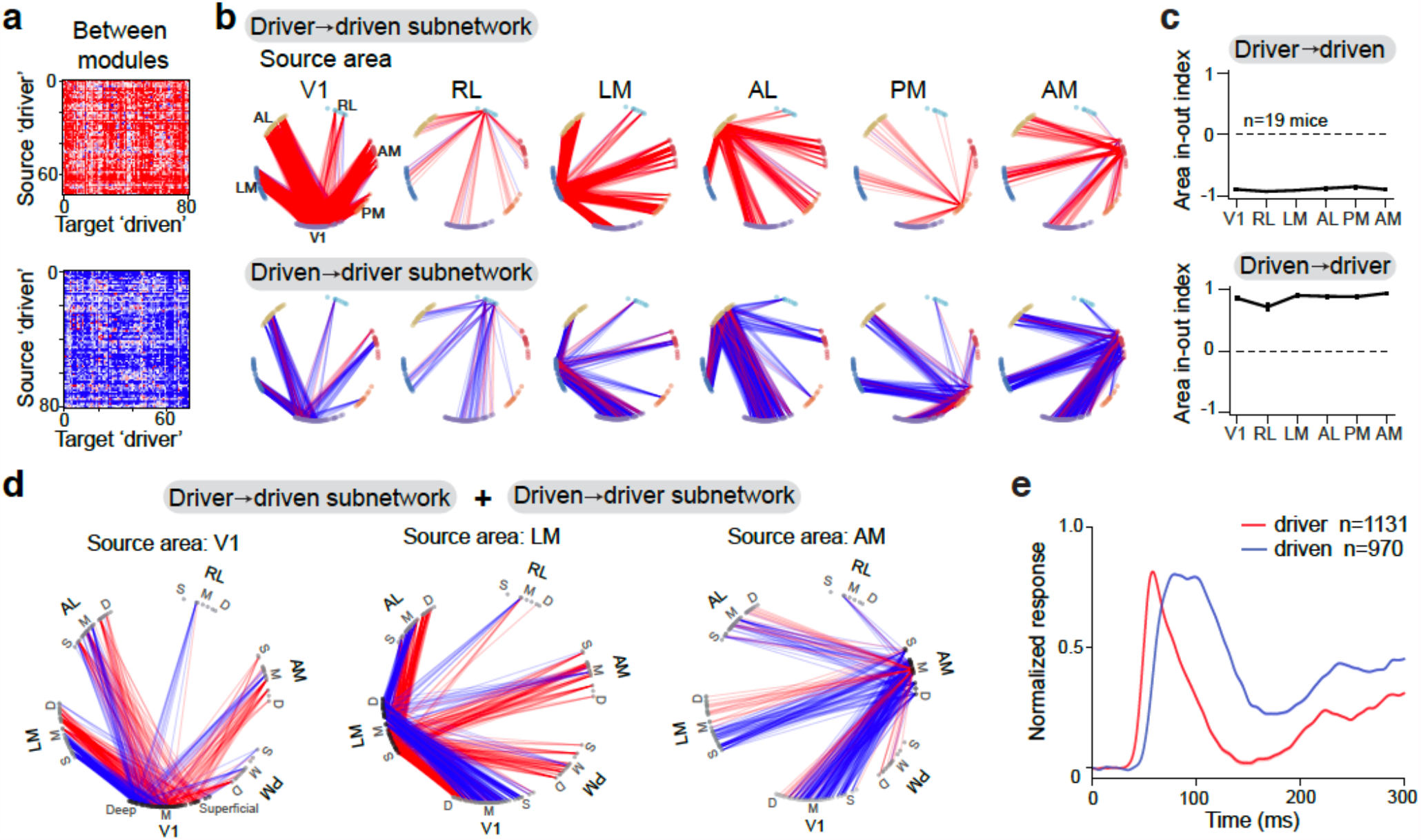
Unidirectional signal flow between distinct modules. **a**) Directed connectivity matrix for between-module subnetworks (*top*: driver (n=74) → driven (n=81); *bottom*: driven → driver) of one example mouse. **b**) Graph representation of strong connections (weight value > 10^−6^) between source area (labeled at the top) and all other areas for between module subnetwork in a). Each node in the graph represents a recording site in cortex. Identity of visual areas is indicated by node color and arranged according to their anatomical location. Nodes in each area are arranged clockwise from superficial to deep layers. Line color indicates the directionality of connection (red: outward projections; blue: inward projections). **c**) Area in-out index averaged across mice (n=19 mice) for between-module subnetworks. **d**) Graph representation of overlaid between-module subnetworks (driver-to-driven and driven-to-driver) for source areas V1, LM and AM. Left: Directed graph representation for between-module subnetworks from source area V1 of one example mouse. Graph representation of strong connections (weight value > 2*10^−6^ for visualization purpose). Line color indicates the directionality of connection. *Red*: output from driver neurons in V1 → driven neurons in other areas; *blue*: input from driver neurons in other areas → driven neurons in V1. There is a clear separation of input and output from source area V1 and the separation is cortical depth dependent. Middle: Source area LM. Right: Source area AM. **e**) Population average of normalized PSTH for the two modules across all areas and all animals. Error bar in this figure represents s.e.m, but it is too small to be visible.

**Figure 6.**
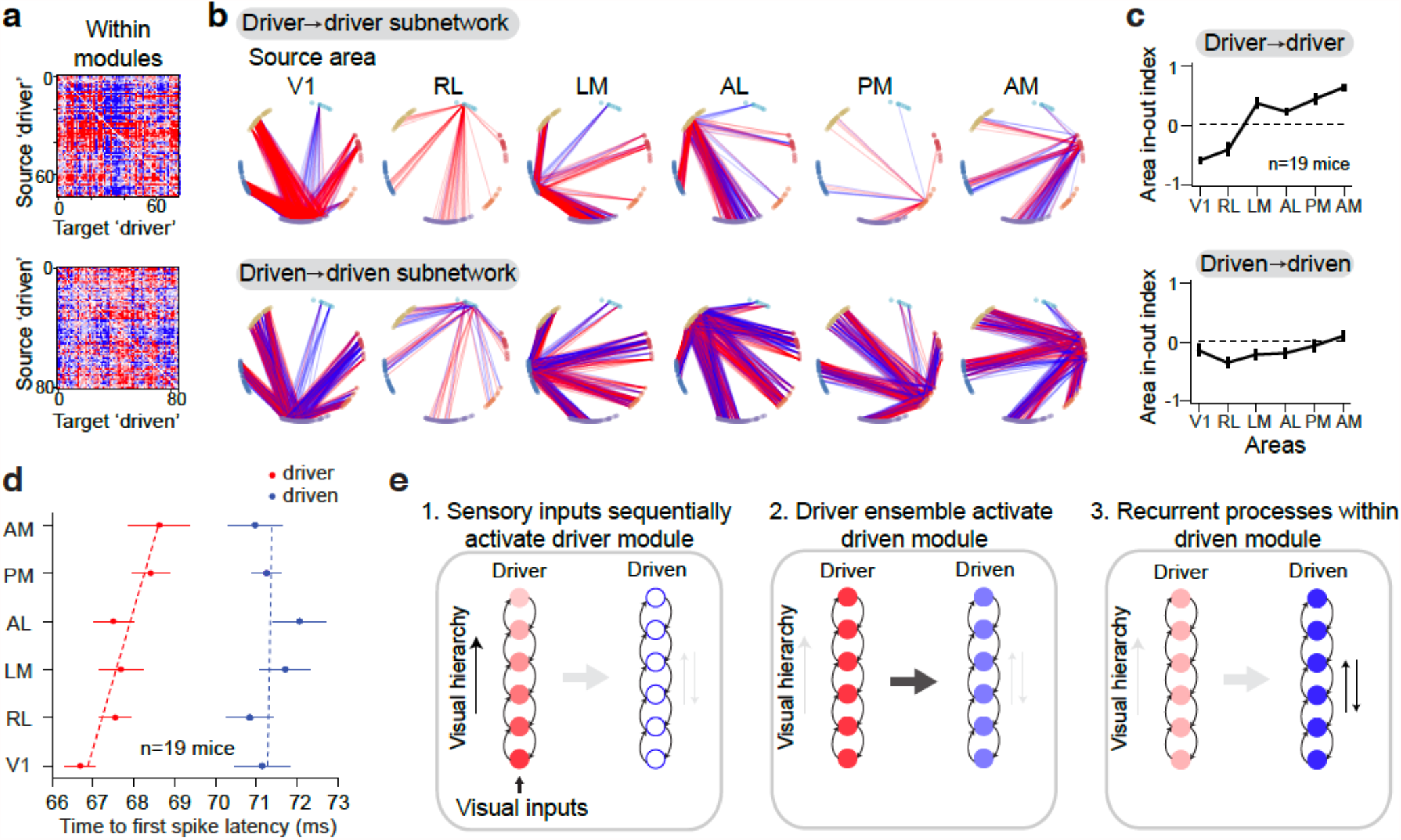
Signal flow within each module indicates a separation of feedforward and recurrent processes. **a**) Directed connectivity matrix for within-module subnetworks (*top*: driver → driver (n=74); *bottom*: driven → driven (n=81)) of the same example mouse as in Fig.4. **b**) Graph representation of strong connections (weight value > 10^−6^) between source area (labeled at the top) and all other areas for within module subnetwork in a). **c**) In-out index averaged across mice (n=19 mice) for within-module subnetworks. **d**) Response latency (time-to-first spike) for different modules across areas (n = 19 mice). **e**) Diagram of the three temporal stages of information flow revealed by data. Intensity of color indicates activation level (darker is stronger). Error bar in this figure represents s.e.m.

To quantify this signal flow from the perspective of individual cortical areas, we define a metric called the ‘area in- out index’, which describes the relative fraction of input versus output connections from a source area for a given subnetwork. An area in-out index of 1 indicates all connections are inputs to the source area, and an index of -1 indicates all connections are outputs (see Methods). For all areas of the driver-to-driven subnetwork the in-out index was close to -1 (Fig. 5c, top), indicating virtually all connections are outputs. In contrast, the in-out index was close to +1 in each area for the driven-to-driver subnetwork across mice (Fig. 5c, bottom). Overlaying these subnetworks for each source area showed a clear separation of inward and outward projections across the cortical depth for each source area (Fig. 5d), with outward projections concentrated in middle to superficial layers of the source area while inward projections concentrated in deep layers. This is consistent with the bias of driver neurons to reside in superficial layers, while driven neurons are biased toward deep layers (Fig. 3b).

The directional communication from driver to driven modules suggests these two modules might be sequentially activated during visual stimulation. To test this, we quantified the stimulus response latency for each module and found that spikes in the driver module preceded the driven module (time to peak = 60.5 ± 0.3 ms versus 80.0 ± 0.6 ms; Rank sum test statistics = -29, p = 7.1e-186) (Fig. 5e). We performed simulations demonstrating that the existence of brief timescale correlations between neurons in these modules does not by necessity entail a temporal offset between modules in the stimulus-triggered average response, and vice versa (Fig. S5). Thus, the directional communication implied by the subnetworks of the adjacency matrix is supported by the temporal progression of signals between the modules.

### Signal transmission within modules

Whereas connectivity between modules was largely unidirectional, the connections *within* each module contained both inputs and outputs indicating more possibilities for recurrent processing (Fig. 6a,b). To explore this within-module communication structure, we analyze the within-module subnetworks (Fig. 6c). Interestingly, for the driver module, the area in-out index systematically increased across the hierarchy: V1 had a negative in-out index and mostly made output connections to driver neurons in other areas; in contrast, driver neurons in AM received more inputs compared to outputs as indicated by a positive in-out index (Fig. 6c, top; Spearman’s correlation with hierarchy is -0.94, p = 0.0048). Within the driven module, connections were more balanced: the in-out index was close to 0 for each area and did not significantly correlate with the hierarchy (Spearman’s correlation, p = 0.16; Fig. 6c bottom).

Theses within-module patterns of connectivity suggest that the driver module neurons relay feedforward signals to driver neurons in other areas along the hierarchy, whereas bi-directional connections within the driven module are positioned to mediate recurrent interactions between areas. Consistent with this, the mean visually evoked response latency within the driver module systematically increased across area hierarchy (Fig. 6d ; correlation with mouse anatomical hierarchy score from (Harris et al., 2019): Spearman’s r = 0.83, p = 0.04). In contrast, visual latencies of neurons in the driven module were delayed relative to the driver module, and did not show an organized progression across the hierarchy (Spearman’s r = 0.14, p = 0.79).

Together, these results suggest a working model in which these two modules support distinct stages of signal propagation: one module transmits feedforward signals about external stimuli along the hierarchy; the other integrates and processes recurrent signals given inputs from the driver module (summarized in Fig. 6e).

### Temporal dispersion of population spiking differs between modules

Modeling studies of feedforward networks have investigated how spiking signals are propagated across modules of sequential processing (Kumar et al., 2010; Reyes, 2003; Vogels and Abbott, 2005). Depending on various network parameters, including connection density and synaptic weights, successive stages can propagate synchronous activity (e.g. a synfire chain; Abeles 1990) or asynchronous fluctuations in firing rate (rate-coded signals). *In vitro* experiments with cultured networks report decreased within-module synchrony of spiking activity as signals are relayed across sequential processing modules (Barral et al., 2019). However, evidence from networks in behaving animals is rare (Zandvakili and Kohn, 2015). The distributed functional modules we describe here could represent sequential stages of signal processing. In this context, we compared the onset synchrony of the first stimulus-evoked spike on a trial-by-trial basis for the driver and driven modules (Fig. 7a-c). Neurons in the driver module were more tightly synchronized with each other compared to those in the driven module (Fig. 7c driver 15.1 ± 4.2 ms and driven 16.4 ± 4.0 ms, n = 8480 trials, Student’s two-tailed t-test statistics T = -20.2, *p* = 2.6e-89). In addition, the dispersion of the trial-wise population response within the driver module (Fig. 7e-f) was also more compact (Fig. 7e Student’s two-tailed t-test statistics T = -31.4, p = 3.3e-211, n = 8480 trials across 19 mice), and occurred earlier (Fig. 7f T = -17.4, *p* = 1.4e-67). These results show that the transmission between the driver and driven module is associated with increased temporal spread of spiking and reduced stimulus onset synchronization. This is consistent with the concept of increasing recurrent interactions deeper into the processing chain (Goris et al., 2014; Lu et al., 2001).

**Figure 7.**
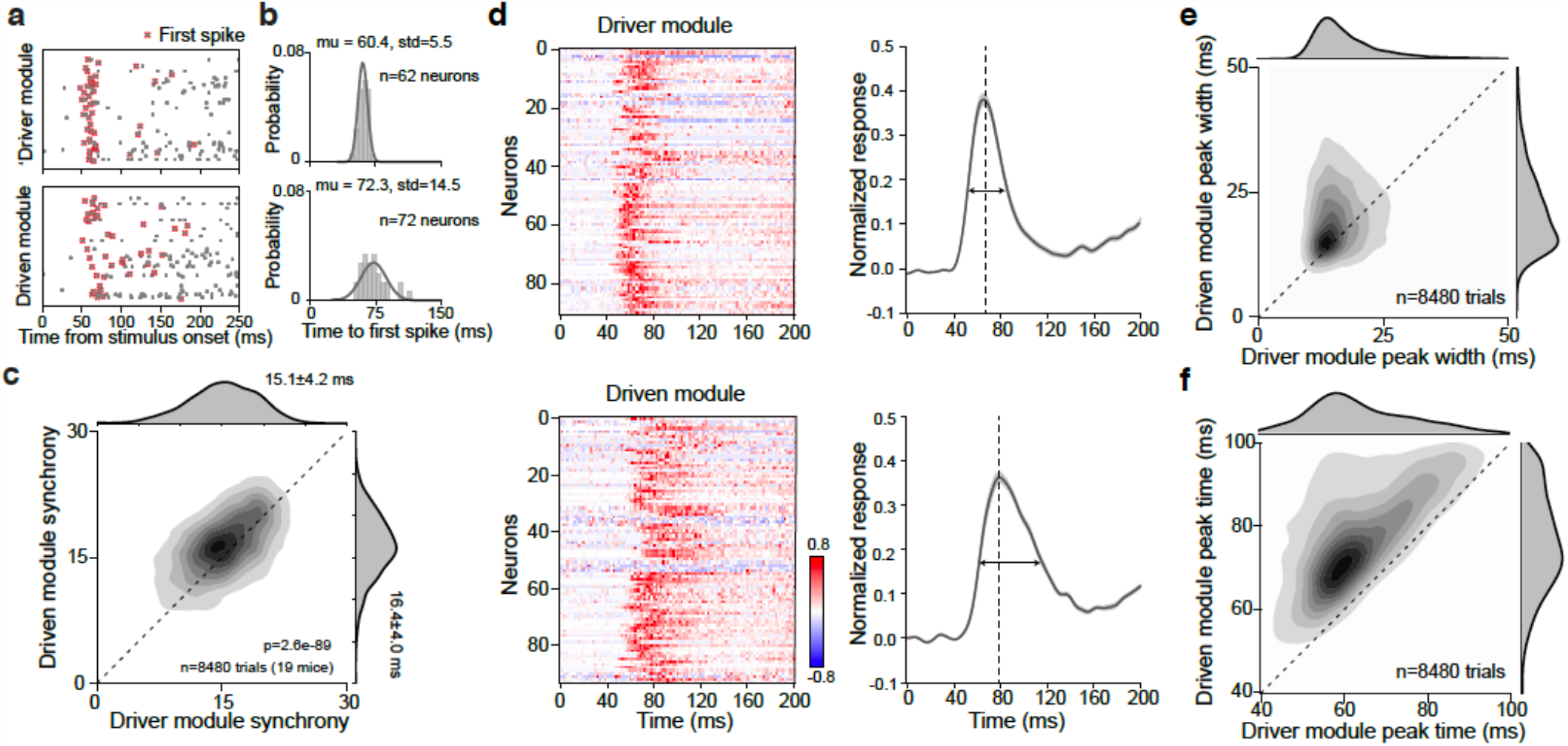
Temporal structure of population spiking differs between modules. **a**) Raster plots for the driver module and driven module of a single trial from an example mouse. Each row is the spike times of one neuron. Red cross highlights the first stimulus-evoked spike. **b**) First spike latency distribution of each module from the trial shown in a). The spread quantifies module onset-latency synchronization in response to stimulus onset. **c**) Joint plot of module onset-latency spread comparing driver and driven module across trials (n = 8480 trials from 19 mice). **d**) Left: normalized PSTHs of individual neurons for driver (n = 91 neurons) and driven (n = 94 neurons) modules from an example mouse. Right: normalized population response of different modules of an example mouse. In this example, the driver module showed a first response peak at 66 ms with spread of 15.24 ms. Driven module showed a first peak at 78 ms with spread of 24.01ms (n = 75 trials). **e**) Joint scatter plot of population response first peak spread across trials and mice. **f**) Joint scatter plot of population response first peak time across trials and mice.

## Discussion

Our results provide a multi-area perspective on signal flow in the mouse visual network. Contrary to a simple feedforward model in which each hierarchical level sequentially transmits signals point-to-point up the chain of areas, we provide evidence for module-based signal propagation that simultaneously involves neurons at multiple levels of the hierarchy. The cluster of neurons we have denoted the *driver* module are engaged earlier in the sensory processing stream, while *driven* neurons are recruited later; this suggests these two modules might represent distinct processing stages. This conclusion is supported by various differences between these two modules including their functional interactions, layer and areas distributions, network convergence/divergence, and within-module temporal coordination.

There are several physiological and computational assumptions we made during experiments and analysis that are important to consider. First, we selected neurons that have an average firing rate larger than a certain threshold (2 spikes/second) when driven by drifting grating stimuli; this constrains our analysis to active units that are stimulus driven. Second, since anatomical connections are retinotopic in the visual cortical areas, to maximize number of measurable functional connections between areas, we selected units with receptive field centers at least 10 degrees away from the edge of the monitor. This selection criterion ensures that the neurons we study have visual receptive field locations that are stimulated by the visual stimulus and facilitated overlap between RFs for functional connectivity estimates. For validation, we did analyze the full functional network of all recorded units before applying any firing rate or RF criterions and found that the units that don’t pass our selection criterions resulted in very low connectivity weights and thus fell into cluster 1, which was dominated by weak connections; thus, these selection criteria are unlikely to influence our conclusions regarding the driver (cluster 2) and driven (clusters 3) modules. Third, when constructing the adjacency matrix from functional connectivity estimated from the jitter-corrected CCG, we did not base our analysis on the shape of the CCG (such as a sharp peak profile, reflecting monosynaptic connections). Instead, we used a permissive definition of directed functional weight (asymmetry of CCG around zero time offset) that likely includes polysynaptic effects. Fourth, we only studied the functional network using drifting grating stimuli (presented at different orientations with many repeats) that does not have a rich spatiotemporal structure that is present in more ecological visual input. This is because drifting gratings stimuli are highly effective in inducing spikes in visual cortex (Engel et al., 1991; Gray and Singer, 1989). To obtain effective connections based on naturalistic stimuli, we will need many more repeats and longer recording times. The influence of using grating stimuli is discussed further below. Finally, our recordings are made in mice that are passively viewing the visual stimuli without being engaged in a task. This likely biases the functional network to reflect bottom-up sensory driven signals rather than top-down signals from higher order cortical regions. Indeed, during spontaneous conditions when no stimulus was shown to the mouse, the connectivity weights between the modules was substantially reduced (Fig. 2), which is consistent with our pervious findings (Siegle et al., 2021). In all, our results and findings should be interpreted with these caveats and constraints in mind.

Despite these caveats, our study reveals a novel organizational principle in the mouse visual cortex in which subsets of neurons distributed across hierarchical levels share functional connectivity and participate in multi-area signal transmission modules. Independent measurements ranging from the composition of simple-like neurons, anatomical distributions and temporal dynamics are all consistent with the different processing stages of these modules, suggesting these multi-area modules are a fundamental organization for visual sensory processing in the mouse. Our general approach can be applied to other brain regions and sensory systems to test the generalization of this organization principle.

### A method for identifying multi-regional functional modules

Community detection methods have been used in human brain imaging studies to identify distributed functional modules at the millimeter scale (Betzel et al., 2018; Power et al., 2011; Sporns and Betzel, 2016), but these techniques have rarely been applied to cellular resolution networks of the cortex (Billeh et al., 2014; Kiani et al., 2015). The clustering algorithm we developed in this study provides a new approach for identifying functional modules in spiking networks and could be helpful for dissecting network substructure in a variety of systems and contexts. Our method for identifying functional modules relied on first quantifying fast timescale functional connectivity (putative synaptic interaction) in the recorded network (Smith and Kohn, 2008a). We define a directed rather than undirected adjacency matrix of interactions between all simultaneously recorded neurons. Our unsupervised clustering method then identifies distinct neuronal populations in the network based on their shared patterns of inferred input and output connectivity. By allowing our algorithm to cluster neurons from all recorded cortical areas together, we uncovered modules that do not map directly onto single cortical areas or layers, although there are area and layer biases. Thus, our approach has the potential for identifying functionally relevant and interacting subpopulations of neurons that are not constrained by standard anatomical parcellation schemes. This is critical, because many behavioral and cognitive operations are likely mediated by such distributed neuronal ensembles (Buzsáki, 2010; Hebb, 1950).

### Separate sets of neurons for distributing and integrating signals

We consistently identified three modules in each mouse based on their shared functional connectivity during visual stimulation. One module is dominated by weak connections, which could be due to non-optimal activation or incompleteness of the recorded network. The other two modules had strong positive connections (driver module) or strong negative connections (driven module) and are the focus of this paper. These modules are determined by their shared functional network connectivity, which is inferred from the asymmetry of the CCGs estimated during drifting gratings. Since drifting gratings are coherent across space and time, they may induce more correlated activity in the population than naturalistic stimuli. However, any boosted correlation should be common to all neurons rather than separate modules. Therefore, the separation of functional modules is unlikely to be simple artifact of the shared inputs. In addition, to minimize the influence of grating stimuli on functional connectivity measurements, we used the jitter-corrected method which largely removes correlations due to shared inputs and slow-timescale co-fluctuations (Smith and Kohn, 2008b) (see Methods and Fig S4). Furthermore, the findings of separate functional modules are consistent with a set of independent measurements, ranging from their anatomical distribution across layers and hierarchy, the temporal dynamics of individual and population of neurons, and the composition of simple and complex-like neurons in each module. These different aspects of evidence are all consistent with the signal flow inferred from the functional connectivity, supporting the conclusion that one module is distributing feedforward sensory signals while the other is responsible for recurrent integration.

Specifically, neurons in the driver module lead activity in the network and show higher divergence implying they could be involved in distributing feedforward signals (Markov et al., 2014b; Sporns, 2013). The driver units also have more transient/faster dynamics compared to the driven units, and their responses are more synchronized (or compact) as a population. Anatomically, the driver units are distributed in the middle and superficial layers and their proportion decreased along the visual hierarchy, consistent with the anatomical distribution of feedforward projecting neurons (Felleman and Van Essen, 1991; Harris et al., 2019; Markov et al., 2014a). These features are consistent with these neurons being more involved in feedforward processing. The driven module, in contrast, is positioned to process recurrent signals. It has higher neuronal convergence, longer sustained response and there are more bi-directional functional connections amongst neurons in this module.

Several theories of cortical processing, including predictive coding (Friston, 2005) and Ullman’s counterstream hypothesis (Shimon Ullman, 1995), posit segregated feedforward and feedback circuits. The functional modules we describe could have implications for these theories. Beyond feedforward and feedback circuits, the idea of separate sets of neurons involved in distributing versus integrating signals has not been sufficiently explored in a sensory processing, where the dominant framework is that of sequential feedforward layers of processing (Riesenhuber and Poggio, 1999). Our findings could have interesting computational implications for the network architecture of cortex. Rather than building a hierarchical networks with sequential stages representing areas, these models can include intercalated groups of neurons that participate in distinct subnetworks with different computational dynamic properties.

The anatomical substrate for the functional modules we identified here is yet to be determined. In the mouse visual cortex, V1 projects to all higher order visual areas (Harris et al., 2019; Wang and Burkhalter, 2007), and single neurons can make branched projections to multiple areas (Han et al., 2018). If these projections were module-specific, this anatomical connectivity could underlie the functional network structure we observe. There is evidence that feedforward and feedback circuits are mediated by separate sets of neurons (Berezovskii et al., 2011; Markov et al., 2014b) and that the rules of local connectivity between V1 neurons depend on the areas to which they project (Kim et al., 2018); these observations of anatomical compartmentalization could help explain the division between the driver and driven modules.

Our observations of the convergence and divergence properties of individual driver and driven neurons may have implications for the cellular anatomy of their axonal projections and dendritic inputs. The higher divergence of the driver neurons suggests that, all else being equal, these neurons could have axons with a greater spatial extent compared to the driven neurons. Similarly, the higher convergence onto driver neurons suggests that these cells may have more spatially extensive dendritic trees. Ultimately, causal perturbations will be necessary to confirm the existence of these two modules and to work out their exact mechanisms of signal transmission. Because the functional modules we describe are highly distributed across different cortical areas and layers, precise labeling and manipulation of neurons in the modules will be technically challenging. Cell type and layer-specific genetic tools such as Cre lines are currently insufficient to study these modules. Photo-stimulation approaches that allow random access activation of both distributed and intermingled cells could be useful for this purpose (Marshel et al., 2019). This method would require that the driver and driven modules be identified prior to photo-stimulation. In principle this could be done using large-scale 2-photon calcium imaging to image activity in distributed populations of neurons across visual cortical areas.

### Future directions

Anatomical connectivity is the scaffold that constrains the observed functional modules, but there are elaborate anatomical projections within and between brain regions that could give rise to different functional network configurations. Future studies can use more complex stimuli to probe how modular network structures depends on different types of inputs, especially naturalistic visual stimuli such as spatiotemporal movies and stimulus visuomotor feedback (e.g. virtual reality) (Saleem et al., 2018). In addition, behavioral states and cognitive task conditions might also restructure the functional networks in the mouse visual cortex. It remains to be determined whether such top-down signals take advantage of the bottom-up functional structure we observe here, perhaps by modulating the gain of transmission between the driver and driven modules. Or whether top-down signals might reconfigure the architecture of interactions to engage separate subsets of neurons than those comprising the driver and driven modules we observe during passive bottom-up stimulation.

Finally, the anatomical hierarchy continues well beyond visual cortex (Harris et al., 2019). More extensive sampling of this network will be necessary to determine the extent to which driver and driven modules extend throughout the cortical and, indeed, the cortico-thalamic system of the mouse or to the brain of other species.

## Acknowledgments

We thank the Allen Institute founder, Paul G. Allen, for his vision, encouragement and support. We thank the Transgenic Colony Management for mouse breeding and Laboratory Animal Services for mouse import and wellness care. We thank the Neurosurgery and Behavior Team for surgical procedures and habituation. We thank Shiella Caldejon for running intrinsic signal imaging experiments, and Rusty Nicovich and Kiet Ngo for collecting optical projection tomography data. We thank the following for helpful discussions: Yazan Billeh, Uygar Sumbul, and Daniel Denman. We thank the following for helpful feedback on manuscript: Daniel Denman, Hannah Choi, Marina Garrett, Gabe Ocker, and Adam Kohn.

## Author contributions

Conceptualization: X.J., J.H.S. and S.R.O.

Investigation, validation and methodology: X.J., J.H.S., S.D., G.H and T.R. Formal analyses: X.J.

Visualization: X.J., S.R.O.

Original draft written by X.J. and S.R.O. with input and editing from J.H.S. and C.K. All co-authors reviewed the manuscript.

## Materials and Methods

### Mice

Mice were maintained in the Allen Institute animal facility and used in accordance with protocols approved by the Allen Institute’s Institutional Animal Care and Use Committee. Four mouse genotypes were used: wild-type C57BL/6J (Jackson Laboratories) (*n* = 11) or Pvalb-IRES-Cre (*n* = 1), Vip-IRES-Cre (*n* = 2), and Sst-IRES-Cre (*n* = 5) mice bred in-house and crossed with an Ai32 channelrhodopsin reporter line. Following surgery, all mice were single-housed and maintained on a reverse 12-hour light cycle. All experiments were performed during the dark cycle.

### Data collection

Experimental data collection followed the procedures described in Siegle, Jia et al., 2019 (Siegle et al., 2019). A summary of these methods is provided below. 13 out of 19 of datasets in this study were previously released on the Allen Institute website via the AllenSDK (https://github.com/AllenInstitute/AllenSDK). The raw data for the rest mice will be available on DANDI drive.

### Surgical methods

All surgical methods used here are the same as (Siegle et al., 2019). Briefly, to enable co-registration across the surgical, intrinsic signal imaging, and electrophysiology rigs, each animal was implanted with a titanium headframe that provides access to the brain via a cranial window and permits head fixation in a reproducible configuration. To implant the headframe, mice were initially anesthetized with 5% isoflurane (1-3 min) and placed in a stereotaxic frame (Model# 1900, Kopf). Isoflurane levels were maintained at 1.5-2.5% for surgery and body temperature was maintained at 37.5°C. Carprofen was administered for pain management (5-10 mg/kg, S.C.). Atropine was administered to suppress bronchial secretions and regulate heart rhythm (0.02-0.05 mg/kg, S.C.). The headframe was placed on the skull and fixed in place with White C&B Metabond (Parkell). Once the Metabond was dry, the mouse was placed in a custom clamp to position the skull at a rotated angle of 20°, to facilitate the creation of the craniotomy over visual cortex. A circular piece of skull 5 mm in diameter was removed, and a durotomy was performed. The brain was covered by a 5 mm diameter circular glass coverslip, with a 1 mm lip extending over the intact skull. The bottom of the coverslip was coated with a layer of silicone to reduce adhesion to the brain surface. At the end of the procedure, but prior to recovery from anesthesia, the mouse was transferred to a photo-documentation station to capture a spatially registered image of the cranial window.

On the day of recording (at least four weeks after the initial surgery), the cranial coverslip was removed and replaced with an insertion window containing holes aligned to six cortical visual areas. First, the mouse was anesthetized with isoflurane (3%–5% induction and 1.5% maintenance, 100% O_2_) and eyes were protected with ocular lubricant (I Drop, VetPLUS). Body temperature was maintained at 37.5°C (TC-1000 temperature controller, CWE, Incorporated). The cranial window was gently removed to expose the brain. An insertion window with holes for probe penetration based on each mouse’s individual visual area map was then placed in the headframe well and sealed with Metabond. An agarose mixture was injected underneath the window and allowed to solidify. The mixture consisted of 0.4 g high EEO Agarose (Sigma-Aldrich), 0.42 g Certified Low-Melt Agarose (Bio Rad), and 20.5 mL ACSF (135.0 mM NaCl, 5.4 mM KCl, 1.0 mM MgCl2, 1.8 mM CaCl2, 5.0 mM HEPES). This mixture was optimized to be firm enough to stabilize the brain with minimal probe drift, but pliable enough to allow the probes to pass through without bending. A layer of silicone oil (30,000 cSt, Aldrich) was added over the holes in the insertion window to prevent the agarose from drying. A 3D-printed plastic cap was screwed into the headframe well to keep out cage debris. At the end of this procedure, mice were returned to their home cages for 1-2 hours prior to the Neuropixels recording session.

### Intrinsic Signal Imaging

Intrinsic signal imaging was performed approximately 15 days after the initial surgery and 25 days before the experiment. Intrinsic signal imaging was used to obtain retinotopic maps representing the spatial relationship of the visual field (or, in this case, coordinate position on the stimulus monitor) to locations within each cortical area (Fig. 1a). The maps made it possible to delineate functionally defined visual area boundaries in order to target Neuropixels probes to retinotopically defined locations in primary and higher order visual cortical areas (Garrett et al., 2014).

### Habituation

Mice underwent two weeks of habituation in sound-attenuated training boxes containing a headframe holder, running wheel, and stimulus monitor. Each mouse was trained by the same operator throughout the 2-week period. During the first week, the operator gently handled the mice, introduced them to the running wheel, and head-fixed them with progressively longer durations each day. During the second week, mice run freely on the wheel and were exposed to visual stimuli for 10 to 50 min per day. The following week, mice underwent habituation sessions of 75 minutes and 100 minutes on the recording rig, in which they viewed a truncated version of the same stimulus shown during the experiment.

### Electrophysiology Experiments

All neural recordings were carried out with Neuropixels probes (Jun et al., 2017). Each probe contains 960 recording sites, a subset of these (374 for “Neuropixels 3a” or 383 for “Neuropixels 1.0”) can be configured for recording at any given time. The electrodes closest to the tip were always used, providing a maximum of 3.84 mm of tissue coverage. The sites are oriented in a checkerboard pattern on a 70 μm wide x 10 mm long shank. The signals from each recording site are split in hardware into a spike band (30 kHz sampling rate, 500 Hz highpass filter) and an LFP band (2.5 kHz sampling rate, 1000 Hz lowpass filter).

The experimental rig was designed to allow six Neuropixels probes to penetrate the brain approximately perpendicular to the surface of visual cortex (Siegle et al., 2019). Each probe was mounted on a 3-axis micromanipulator (New Scale Technologies, Victor, NY), which were in turn mounted on a solid aluminum plate, known as the probe cartridge. The mouse was placed on the running wheel and fixed to the headframe clamp. The tip of each probe was aligned to target the desired retinotopic region in each area. Brightfield photo-documentation images were taken with the probes fully retracted, after the probes reached the brain surface, and again after the probes were fully inserted. An IR dichroic mirror was placed in front of the right eye to allow an eyetracking camera to operate without interference from the visual stimulus. A black curtain was then lowered over the front of the rig, placing the mice in complete darkness except for the visual stimulus monitor.

Neuropixels data was acquired at 30 kHz (spike band) and 2.5 kHz (LFP band) using the Open Ephys GUI (Siegle et al., 2017). Gain settings of 500x and 250x were used for the spike band and LFP band, respectively. Each probe was either connected to a dedicated FPGA streaming data over Ethernet (Neuropixels 3a) or a PXIe card inside a National Instruments chassis (Neuropixels 1.0). Raw neural data was streamed to a compressed format for archiving which was extracted prior to analysis.

### Cortical Area Targeting

To confirm the identity of the cortical visual areas, images of the probes taken during the experiment were compared to images of the brain surface vasculature taken during the ISI session (see above). Vasculature patterns were used to overlay the visual area map on an image of the brain surface with the probes inserted (Fig 1a). To maximize measurable functional connectivity across areas, we targeted the center of gaze in all areas (except for RL, which targeted the center of mass because of geometry) with overlapping receptive fields (RF) guided by a retinotopic map. Targeting was validated by mapping receptive fields of all sorted units with small Gabor patches presented at different locations on the screen (see below). All analysis was restricted to neurons with well-defined receptive fields within the screen boundaries.

### Visual Stimulus

Visual stimuli were generated using custom scripts based on PsychoPy (Peirce, 2007) and were displayed using an ASUS PA248Q LCD monitor, with 1920 × 1200 pixels (21.93 in wide, 60 Hz refresh rate). Stimuli were presented monocularly, and the monitor was positioned 15 cm from the mouse’s right eye and spanned 120° x 95° of visual space prior to stimulus warping. Each monitor was gamma corrected and had a mean luminance of 50 cd/*m*^2^. To account for the close viewing angle of the mouse, a spherical warping was applied to all stimuli to ensure that the apparent size, speed, and spatial frequency were constant across the monitor as seen from the mouse’s perspective.

#### Visual stimuli for receptive fields (RFs)

Receptive field location was mapped with small Gabor patches. The receptive field mapping stimulus consisting of 2 Hz, 0.04 cycles per degree drifting gratings (3 directions: 0°, 45°, 90°) with a 20° circular mask. These Gabor patches randomly appeared at one of 81 locations on the screen (9 × 9 grid with 10° spacing) for 250 ms at a time, with no blank interval.

#### Visual stimuli for current source density (CSD)

Current source density for layer estimation used the full-field flash stimuli (a series of dark or light full field image with luminance = 100 cd/*m*^2^). lasting 250 ms each and separated by a 1.75 second inter-trial interval.

#### Visual stimuli for functional connectivity

Functional connectivity during the stimulus-driven condition was measured using drifting grating stimuli, which were presented at 4 directions (0°, 45°, 90°, 135°), with temporal frequency equal to 2 cycle/sec and contrast equal to 0.8. In each trial, the grating is presented for 2 sec followed by 1 sec gray screen. Each condition was presented for 75-100 trials.

### Spike Sorting

Prior to spike sorting, the spike-band data passed through 4 steps: DC offset removal, median subtraction, filtering, and whitening. First, the median value of each channel was subtracted to center the signals around zero. Next, the median across channels was subtracted to remove common-mode noise. The median-subtracted data file is the input to the Kilosort2 Matlab package (https://github.com/mouseland/kilosort2), which applies a 150 Hz high-pass filter, followed by whitening in blocks of 32 channels. The filtered, whitened data is saved to a separate file for the spike sorting step.

Kilosort2 was used to identify spike times and assign spikes to individual units (Stringer et al., 2019). Kilosort2 attempts to model the complete dataset as a sum of spike “templates.” The shape and locations of each template is iteratively refined until the data can be accurately reconstructed from a set of *N* templates at *M* spike times, with each individual template scaled by an amplitude, *a*. A critical feature of Kilosort2 is that it allows templates to change their shape over time, to account for the motion of neurons relative to the probe over the course of the experiment. Stabilizing the brain using an agarose-filled plastic window has virtually eliminated probe motion associated with animal running, but slow drift of the probe over ∼3-hour experiments is still observed. Kilosort2 is able to accurately track units as they move along the probe axis, eliminating the need for the manual merging step that was required with the original version of Kilosort (Pachitariu et al., 2016). The spike-sorting step runs in approximately real time (∼3 hours per session) using a dual-processor Intel 4-core, 2.6 GHz workstation with an NVIDIA GTX 1070 GPU. We used the default parameters in Kilosort2, with an initial threshold of 12, and a final-pass threshold of 8.

The Kilosort2 algorithm will occasionally fit a template to the residual left behind after another template has been subtracted from the original data, resulting in double-counted spikes. This can create the appearance of an artificially high number of ISI violations for one unit or artificially high zero-time-lag synchrony between nearby units. To eliminate the possibility that this artificial synchrony will contaminate data analysis, the outputs of Kilosort2 are post-processed to remove spikes with peak times within 5 samples (0.16 ms) and peak waveforms within 5 channels (∼50 microns).

Kilosort2 generates templates of a fixed length (2 ms) that matches the time course of an extracellularly detected spike waveform. However, there are no constraints on template shape, which means that the algorithm often fits templates to voltage fluctuations with characteristics that could not physically result from the current flow associated with an action potential. The units associated with these templates are considered “noise,” and are automatically filtered out based on 3 criteria: spread (single channel, or >25 channels), shape (no peak and trough, based on wavelet decomposition), or multiple spatial peaks (waveforms are non-localized along the probe axis).

Following the spike sorting step, data for each session was uploaded to the Allen Institute Laboratory Information Management System (LIMS). Each dataset was run through the same series of processing steps using a set of project-specific workflows (AllenSDK v1.0.2) in order to generate NeurodataWithoutBorders (NWB) files used for further analysis.

## Analysis Methods

### Dataset

In total, units from 19 mice were included in our functional connectivity analysis. Spike sorting, quality control, and preprocessing steps followed the same procedures as (Siegle et al., 2019). 13 out of 19 of these datasets were previously released on the Allen Institute website via the AllenSDK (https://github.com/AllenInstitute/AllenSDK). On average, 632 ± 18 sorted cortical units were simultaneously recorded in each mouse. We set a firing rate threshold to select units for functional connectivity analysis. Firing rate (FR) was defined as the average number of spikes in a window from 50 ms to 500 ms after the onset of the drifting gratings stimulus. Only units with mean FR > 2 spikes/second were used for pairwise cross-correlogram (CCG) calculation, which resulted in an average of 356 ± 7 units in each mouse (*n* = 6773 units in total). Because functional connectivity varies with receptive field position (Jia et al., 2013), we further constrained the dataset to include units with receptive field centers at least 10 degree away from the edge of the monitor (see Visual receptive fields section below. After filtering by receptive field location, we ended up with 184 ± 8 per mouse used for the final clustering procedure (*n* = 3487 units in total). After applying clustering on the functional connectivity matrix constrained by both FR and RF location in each mouse, the total numbers of units belonging to each cluster were: n_cluster1 = 1386, n_cluster2 = 1131, n_cluster3 = 970.

### Quantification and statistical analysis

All analyses were performed in Python. The main analysis packages used in this paper are Scipy (Virtanen et al., 2020), scikit-learn (Pedregosa et al., 2011), statsmodels (Seabold and Perktold, 2010), and network (Hagberg et al., 2008). Error bars, unless otherwise specified, were computed as standard error of the mean. When comparing the difference between two independent variables, if their distribution is Gaussian like (normality test), we used Student’s *t*-test; if their distribution is non-Gaussian, we used a rank sum test. When testing whether a distribution is significantly different from 0, we used a one-sample *t*-test. When comparing variables between modules across cortical areas, we used two-way analysis of variance (ANOVA) to assess both the main effect between modules and whether there is any interaction across areas. When comparing similarity to the previously established anatomical visual hierarchy in mouse (Harris et al., 2019), we calculated the correlation between our measured variable (e.g. first spike latency) and the previously calculated hierarchy score (V1: -0.50, RL: -0.12, LM: -0.13, AL: 0.00, PM: 0.11, AM: 0.29), using Spearman’s correlation to estimate the rank order significance. Statistical details and *p*-values can be found in the Results section or figure legends.

### Visual receptive fields

Receptive fields were mapped with Gabor patches (20 degree each; 3 different orientations (0, 45, 90), temporal frequency = 2 cyc/s, spatial frequency = 0.04 cyc/deg) shown randomly at 81 different locations (9 × 9 grid, 10° separation between pixel centers) with gray background on a 120° x 95° monitor (1920 × 1200 pixels, 21.93 inches wide, 60 Hz refresh rate). The receptive field map (RF) for one unit is defined as the mean 2D histogram of spike counts at each of 81 locations, each pixel covers a 10° x 10° square. The receptive field was then thresholded at 20% of maximum response to remove potential noisy pixels. Then, a 2D Gaussian

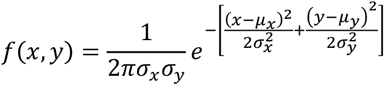

was fit to the thresholded visual receptive map to estimate the center of the receptive field location.

### Peristimulus time histogram (PSTH)

To visualize the temporal dynamics of a neuronal population (Fig. 1, Fig. 3, and Fig. 4), the activity of each neuron was binned at 1 ms, averaged across trials (*n* = 75), smoothed with a Gaussian filter with standard deviation of 3 ms, baseline subtracted (baseline period from 0 to 0.03s relative to stimulus onset), and normalized by dividing the maximum of the response between 0 to 1.5 s after stimulus onset. The normalized PSTHs of individual neuron were averaged within a neuronal population; the error bars indicate standard error of the mean across neurons.

### Functional connectivity

We analyzed functional interactions between pairs of simultaneously recorded neurons by calculating the spike train cross-correlogram (CCG) (Jia et al., 2013; Smith and Kohn, 2008c; Zandvakili and Kohn, 2015). For a pair of neurons with spike train *x*_1_ and *x*_2_, the CCG is defined as:

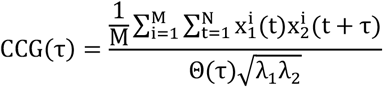

where *M* is the number of trials, *N* is the number of bins in the trial, 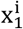 and 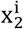 are the spike trains of the two units on trial i, τ is the time lag relative to reference spikes, and λ_1_ and λ_2_ are the mean firing rates of the two units. The CCG is essentially a sliding dot product between two spike trains. θ(τ) is the triangular function which corrects for the overlapping time bins caused by the sliding window. To correct for firing rate dependency, we normalized the CCG by the geometric mean spike rate. An individually normalized CCG is computed separately for each drifting grating orientation and averaged across orientations to obtain the CCG for each pair of units.

The jitter-corrected CCG was created by subtracting the expected value of CCGs produced from a resampled version of the original dataset with spike times randomly perturbed (jittered) within the jitter window (Harrison and Geman, 2009; Smith and Kohn, 2008c). The correction term (*CCG*_*jittered*_) is the true expected value which reflects the average over all possible resamples of the original dataset. *CCG*_*jittered*_ is normalized by the geometric mean rate before subtracting from *CCG*_*original*_. The analytical formula used to create a probability distribution of resampled spikes was provided in Harrison and Geman, 2009. This method disrupted the temporal correlation within the jitter window, while maintaining the number of spikes in each jitter window and the shape of the PSTH averaged across trials.

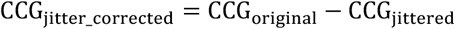

For our measurement, a 25 ms jitter window was chosen based on previous studies (Jia et al., 2013; Zandvakili and Kohn, 2015). This jitter-correction method removes both the stimulus-locked component of the response, as well as slow fluctuations larger than the jitter window. The remaining fast timescale correlation is more likely to be related to signal propagation between two neurons. Therefore, the jitter-corrected CCG reflects temporal correlations between a pair of neurons within the jitter-window (25ms).

We then calculated the directed connection weight by subtracting the sum of (−13 to 0) ms of the CCG from the sum of (0 to 13) ms of the jitter-corrected CCG (Fig. 1d). The 13ms window was defined as half of the 25 ms jitter window we used, and also because real functional delay between neurons in mouse occur on the timescale of milliseconds to tens of milliseconds (Siegle et al., 2019). The resulting value indicates the strength and the sign indicates the directionality of the functional connection between a pair of neurons. Computing this for all pairs of neurons produced a directional, cellular-resolution connectivity matrix for each mouse (Fig. 1e).

### Clustering

#### Non-randomness

We first tested whether there is modular structure (non-randomness) in the measured connectivity matrix by computing the graph spectrum (based on spectral graph theory (Spielman, 2008). The eigenvalues of a graph are defined as the eigenvalues of its adjacency matrix (Fornito et al., 2016). The set of eigenvalues of a graph forms a graph spectrum. The randomness of the matrix is quantified by comparing the graph spectrum of the original connectivity matrix with its shuffled connectivity matrix, where the x and y axis are shuffled independently, and a randomly generated connectivity matrix with the same size. We found that the graph spectrum of the original matrix showed significantly higher explained variance by the top eigenvalues than the shuffled matrix and the random matrix, suggesting that the measured connectivity matrix has non-random structure (Fig. S1).

#### Defining the number of clusters

The number of clusters was determined using several complementary methods (Fig. S3a):

1. The Elbow method estimates the percentage of variance explained for a given number of *k*. The number of cluster 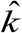 is estimated at the point when the curve turns into a plateau. The following measure represents the sum of within-cluster distances (pairwise distances) between all points in a given cluster *C*_*k*_ containing *n*_*k*_ points:

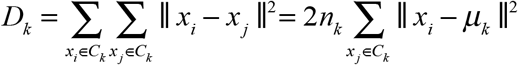 Adding the normalized within-cluster sum-of-squares gives a measure of the compactness of our clustering, or the pooled within-cluster sum of squares around the cluster means:

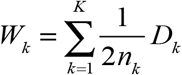

*W*_*k*_ increases monotonically with number of clusters *k*. The number of clusters is chosen at the point where the marginal gain drops (or the point slope change most dramatically), the ‘elbow’.
2. Gap statistics (Tibshirani et al., 2001) seeks to standardize the comparison of log*W*_*k*_ with a null reference distribution of the data, i.e. a distribution with no obvious clustering. The estimate for the optimal number of clusters *K* is the value for which log*W*_*k*_ falls the farthest below this reference curve. This information is contained in the following formula for the gap statistic:

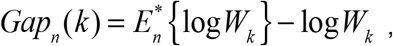 Where 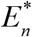 denotes the expectation under a sample of size *n* from the reference distribution. The estimate 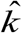 will be the value maximizing *Gap*_*n*_ (*k*) after we take the sampling distribution into account.
3. Clustering density estimates the data distribution density for a given *k* by calculating a density function *f* (*k*) (Pham et al., 2005). The value of *f* (*k*) is the ratio of the real distortion to the estimated distortion. When the data are uniformly distributed, the value of *f* (*k*) is 1. When there are areas of concentration in the data distribution, the value of *f* (*k*) decreases. Therefore, the number of 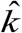 clusters is determined by finding the minimum value of *f* (*k*).

Combining the estimation of 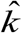 using the above three methods, we determined the optimal number of clusters to be 3.

#### Method for clustering

In order to find neurons that have correlated connectivity patterns to the rest of the network, we clustered the directed connectivity matrix by treating the connectivity pattern from each source neuron to all target neurons as features (Fig. 1f and Fig. S2). To reduce noise, we projected the connectivity features into a lower dimensional space with principal component analysis (PCA), only keeping the top principal components that explained 80% of total variance. We then applied a consensus clustering method (Monti et al., 2003) with k-means to obtain robust clusters that are not biased by random initial conditions. First, we constructed a co-clustering association matrix by running *k*-means with different initial conditions 100 times (reached stable co-clustering). Each entry in the matrix represents the probability of two units belonging to the same cluster. Then, we clustered the association matrix with hierarchical clustering to determine the cluster labels. The number of clusters was determined using methods described in the previous section.

#### Comparing different clustering methods

Our consensus clustering was based on *k*-means clustering methods, which measures the compactness of points based on features in the reduced PCA space (see above). We compared this clustering method with two other clustering methods to detect modular structure in the adjacency matrix: the spectral clustering method (sklearn.cluster.SpectralClustering) and bi-clustering method (sklearn.cluster.SpectralBiclustering).

Spectral clustering determines the clusters based on the connectivity of data points: points that are connected or immediately next to each other are placed in the same cluster. In spectral clustering, the data points are treated as nodes of a graph, and the clustering is treated as a graph partitioning problem. The nodes are then mapped to a low-dimensional space that can be easily segregated to form clusters. The spectral clustering is carried out in 3 steps: 1. Compute a similarity graph (*k*-nearest neighbors). 2. Project the data onto a low-dimensional space (compute Graph Laplacian, and eigenvalues and eigenvector for L). 3. Create clusters (based on the eigenvector corresponding to the 2^nd^ eigenvalue to assign values to each node, then split the nodes with *k*-means for the given number of clusters).

Biclustering (or block clustering) is a method to simultaneously cluster the rows and columns of a matrix. For a *m* (sample) by *n* (feature) matrix, the algorithm generates biclusters, which are a subset of rows that exhibit similar connectivity pattern across a subset of columns.

The results of consensus clustering, spectral clustering, and biclustering of the functional connectivity matrix are shown in (Fig. S3c). The three methods showed relatively consistent clustering results (Fig. S3d) in detecting units that belong to the three clusters dominated by different weight pattern. Therefore, our clustering findings are general and do not depend on the specific clustering method we used.

#### Cluster quality

We used *two methods*, which were previously used to evaluate spike sorting cluster quality (Siegle et al., 2019), to quantify neuronal population cluster quality given different number of clusters (Fig. S3b). The *d*-prime (*d’*) was calculated using Fisher’s linear discriminant analysis to find the line of maximum separation in PC space (Hill et al., 2011). d’ indicates the unbiased separability of the cluster of interest from all other clusters. The higher the value, the more distinguishable are the clusters. Hit-rate was calculated with nearest-neighbors method (n_neighbors = 3), which is a non-parametric estimate of exemplar contamination in each cluster. For each unit belonging to the cluster of interest, the three nearest units in principal-component space are identified. The “hit rate” is defined as the fraction of these units that belong to the cluster of interest. This metric is based on the “isolation” metric from (Chung et al., 2017). The higher the value, the less contamination in each cluster.

### Module distribution

The area distribution of neurons within each module was quantified by calculating the proportional number of units in one area relative to the total number of units in all areas for a given module. The proportion of one module across areas sums to 1. To minimize sampling bias across areas, we subsampled units in each area to match the number of units across areas. The final result was a bootstrapped mean (sampling with replacement to match the number of units in each area; n_boot = 100). Error bars represents the bootstrapped standard deviation across all units in all mice. Results are only shown for the ‘driver’ and ‘driven’ modules. No systematic area bias was observed for cluster 1 (the cluster with non-significant connection) units (result not shown).

The distribution of each neuronal module across layers was quantified by first dividing units into superficial, middle, and deep layers according to the location of layer 4 estimated from the CSD. We then calculating the proportion of units across these three layers for a given neuronal module. To minimize sampling bias across layers, we subsampled units in each layer to match the number of units across layers. Means and error bars were calculated using the same bootstrapping method as for the area distributions.

### Graph creation

To create graphs visualizations (Fig. 5b; 6b), we first condensed our single-unit connectivity matrix to a single-recording-site connectivity matrix by combining units with peak channels on the same electrode. Then, we treat each site as a node in the graph. For an intuitive representation, nodes belonging to the same cortical area are close by and arranged clockwise from superficial to deep layer. The location of each area is determined by the top-down view of the physical locations of visual areas on the left hemisphere (Fig. 1a). The edges of the graph represent connections between sites, with red lines indicating projections from the source unit (positive weight) and blue lines indicating projections back to the source unit (negative weight). The threshold for significant connections is defined as an absolute weight larger than 10^−6^ coincidences/spike, which is about half of the standard deviation of the weight distribution across all mice (Fig. S4a).

### Divergence and convergence degree

Divergence degree is similar in concept to the outdegree of a graph. It is defined as the proportion of significant positive connections (weight > 10^−6^) from a source neuron to the rest of the network (*N* neurons). *C*_*i*,+_ represents the number of positive connections from neuron *i* to the network.

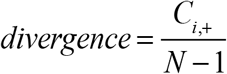

Convergence degree is similar in concept to the indegree of a graph. It is defined as the proportion of significant negative connections (weight < -10^−6^) to source neuron *i* from the rest of the network.

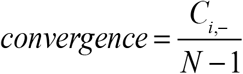

### Temporal dynamics analysis

#### Response latency

Two different measurements were used to estimate response latency. The peak response latency was defined as the time when a neuron’s response reached its first peak after stimulus onset. The time to first spike was estimated in each trial by looking for the time of the first spike 30 ms after stimulus onset. If no spike was detected within 250 ms after stimulus onset, that trial was not included. The overall latency for each unit was defined as the mean time to first spike across trials.

#### Population onset response synchrony

We used the spread of time-to-first spike for all neurons within a module for a single trial as an indicator of population response onset synchronization. The spread was calculated by fitting a Gaussian to each trial’s time-to-first-spike distribution:

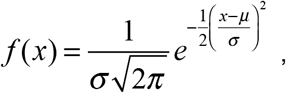

where x is the spike time relative to stimulus onset, *µ* is an estimate of the average time-to-first-spike and *σ*is an estimate of the spread of the time-to-first spike distribution for one trial.

#### Population response spread of the first peak

To quantify spike the response spread of a neuronal ensemble, we estimated the width of the population PSTH for each trial. The PSTH was calculated with 2 ms bins, and convolved with a Gaussian kernel of width 5 ms. The properties of the first peak were estimated using scipy.signal.find_peaks. The peak width represents the half-width at half maximum of the peak, while peak height represents the maximum of the peak. The spike spread of a neuronal population is an important parameter for quantifying how signals are transmitted through a feedforward network.

### Area in-out index

To quantify the proportion of projections out from a source area relative to inputs back into the source area, we defined the area in-out index as

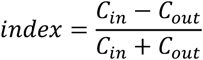

where *C*_*out*_ is the number of connections from source to other areas in the given network and *C*_*in*_ is the number of connections from other areas to source area. This index reflects the asymmetry of in-and-out degree of a source area. When the value is close to -1, the source area is dominated by outward projections (positive weights). When the value is close to 1, the source area is dominated by inward projections (negative weights). When the value is 0, the source area has balanced outward and inward connections.

### Layer definition

We estimated the depth of the middle layer of cortex by first calculating the current source density (CSD) using simultaneously recorded local field potentials (LFP). The CSD was computed using the method in (Stoelzel et al., 2009), using the LFP within 250 ms after stimulus onset. First, we calculated the average evoked (stimulus locked) local field potential at each recording site. Next, we duplicated the uppermost and lowermost field traces and smoothed these signals across sites

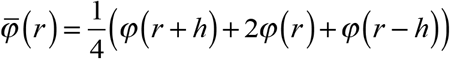

where *φ* is the field potentials, *r* is the coordinate perpendicular to the layers, *h* is the spatial sampling interval (40 μm in our case). Then, we calculated the second spatial derivative

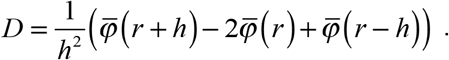

In the resulting CSD map, current sinks are indicated by downward deflections and sources by upward deflections. To facilitate visualization, we smoothed the CSD with 2D Gaussian kernels (*σ*_*x*_ = 1; *σ*_*x*_ = 2). To find the middle layer, we defined the first sink within 100 ms after stimulus onset as the input layer (center channel) by searching for the local maximum on the CSD map (first sink), followed by source.

We used the middle layer estimation for the calculation of layer distribution bias of ‘driver’ and ‘driven’ modules. We partitioned the cortical layers into three layers: middle layer (center channel ± 8 channels, which is ± 40μm), superficial layer (channels above middle layer), and deep layers (channels below middle layer and above white matter).

### Simulations to test mathematical relationship between PSTH shape and CCG sharp peak

Because we observed that the functional connectivity defined ‘driver’ module responded earlier than the ‘driven’ module (Fig. 5e), we wondered whether the brief timescale relationship of ‘driver’ leading ‘driven’ was a consequence of the general latency reflected in averaged PSTH. Even though our jitter-correction method should have removed stimulus-locked components and the observed directionality should only reflect brief-timescale signal transmission, we still wanted to rule out the possibility that the observed asymmetry in the CCG is merely a reflection of the trial-averaged PSTH latency.

We used a simple simulation to carry out positive and negative controls (Fig. S5). The negative control tested whether two neurons with correlated, but temporally offset, PSTH traces will necessarily show significant peaks in their jitter-corrected CCG (25 ms jitter window). The positive control tested whether two neurons with uncorrelated PSTH traces can produce a significant peak in their CCG if we artificially introduce millisecond-timescale correlations. The mathematical expression of the tests is formulated as follows:

Given two PSTH traces: *λ*_1_(*t*) and *λ*_2_(*t*), we simulated *Poisson* spike trains:

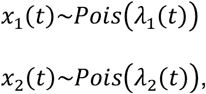

over time *T* for 100 repeats, where the PSTHs of the two simulated spike trains (*X*_1_ and *X*_2_) matched the shape of *λ*_1_(*t*) and *λ*_2_(*t*). Synchronized spikes were introduced to *x*_1_(*t*) and *x*_2_(*t*) only for the positive control. CCGs before and after jitter correction were calculated between *X*_1_ and *X*_2_. We found that brief timescale correlations between two neurons (identified by significant peaks in the CCG) do not depend on the shape and relative timing of their PSTHs, but—as expected—reflect only their fine-timescale temporal relationship.

## Data and code availability

The majority of the data in this study (13 of 19 experiments) was publicly released as an open dataset on the Allen Institute website in October 2019, and is available via the AllenSDK (https://allensdk.readthedocs.io/en/latest/visual_coding_neuropixels.html). Additional data and software will be deposited to Github.

**Supplemental Figure 1.**
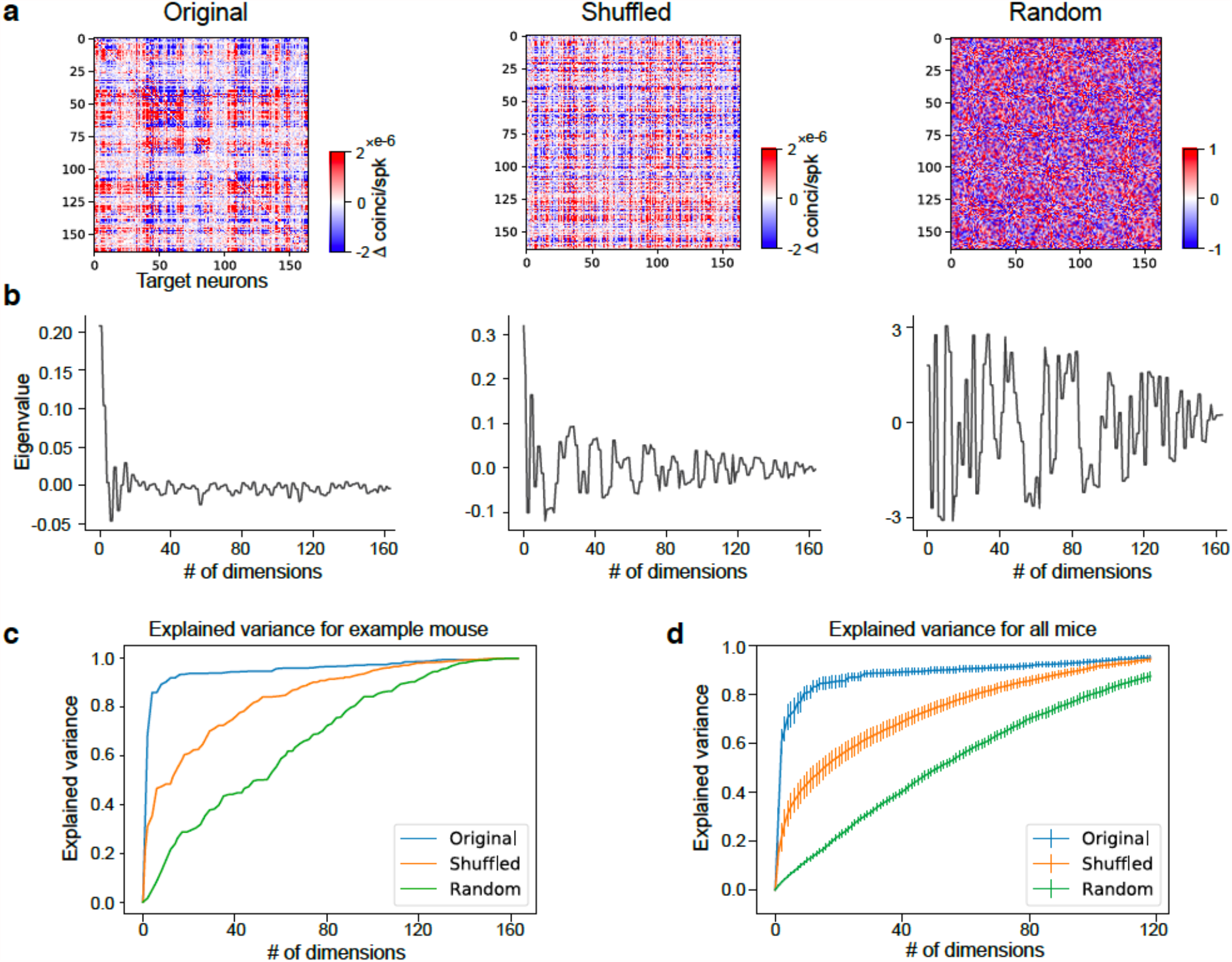
Non-randomness of the functional connectivity matrix. **a**) Functional connectivity matrix from one example experiment (units are sorted by depth and areas). The shuffled matrix control was generated by randomly shuffling the x and y axes of the original matrix. The random matrix control was generated by simulating pure random connections between units. **b**) Eigenvalues of the three matrices in a). **c**) Explained variance based on eigenvalues in b). The original connectivity matrix showed the highest explained variance with relative few dimensions suggesting the presence of organized structure in the matrix compared to the shuffled and random controls. **d**) Quantification of explained variance for the measured and control connectivity matrices across mice (n=19 mice). Error bars indicate standard error of the mean.

**Supplemental Figure 2.**
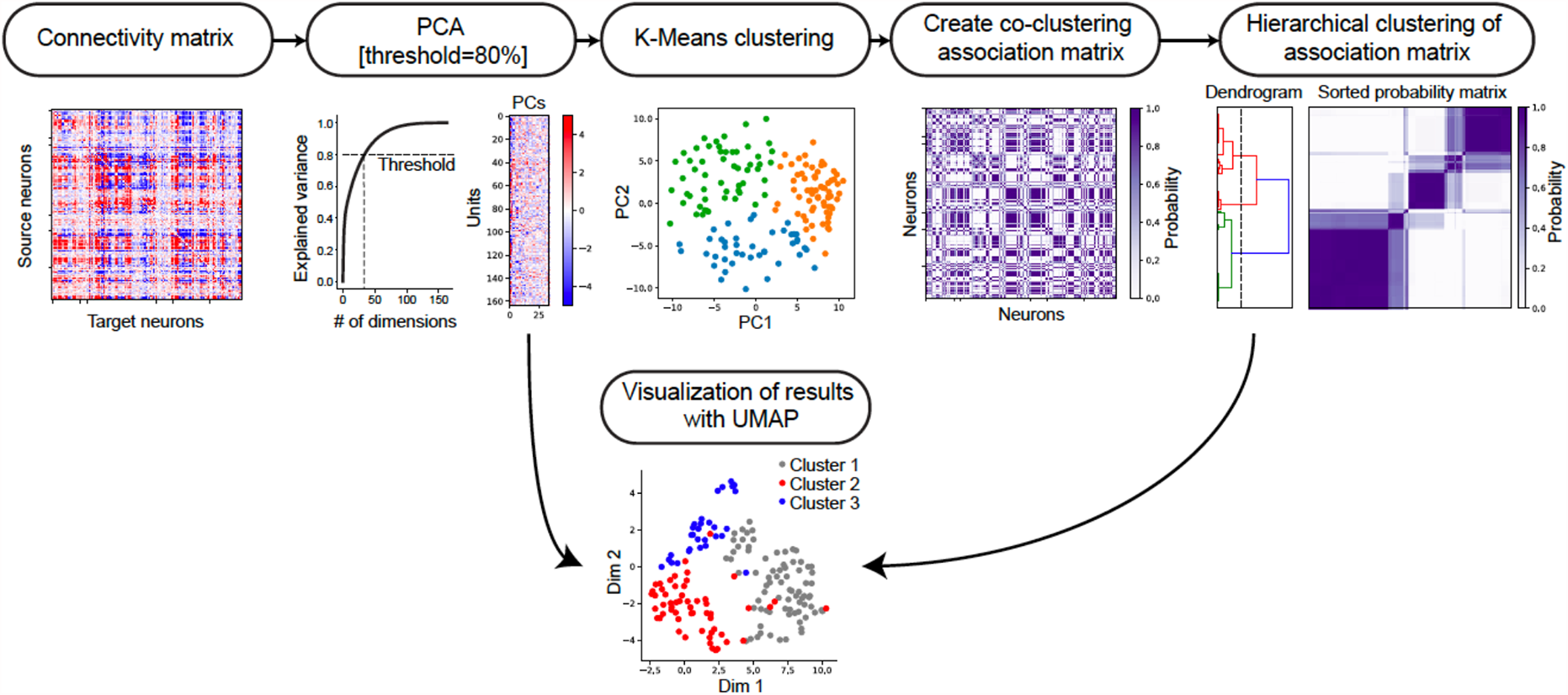
Illustration of clustering method for identifying neuronal modules. Treating the connectivity pattern from each unit to the network as features, we first applied principle component analysis to reduce noise in the original connectivity matrix by keeping the top principle components that explain 80% of the total variance. Then we applied consensus clustering (Monti et al., 2003) to the PCs, by running K-means using the PCs 100 times until reaching a stable co-clustering association matrix, where each entry represents the probability of two units belonging to the same cluster. Finally, we applied hierarchical clustering to the association matrix and kept 3 clusters as the classification result, which was independently validated using the methods in Fig. S3A. As a result, the units are clustered based on their connectivity pattern into three modules. The bottom UMAP (McInnes et al., 2018) is a visualization of 2D embedding with all the PCs and the layout agrees with the cluster labels from the consensus clustering result.

**Supplemental Figure 3.**
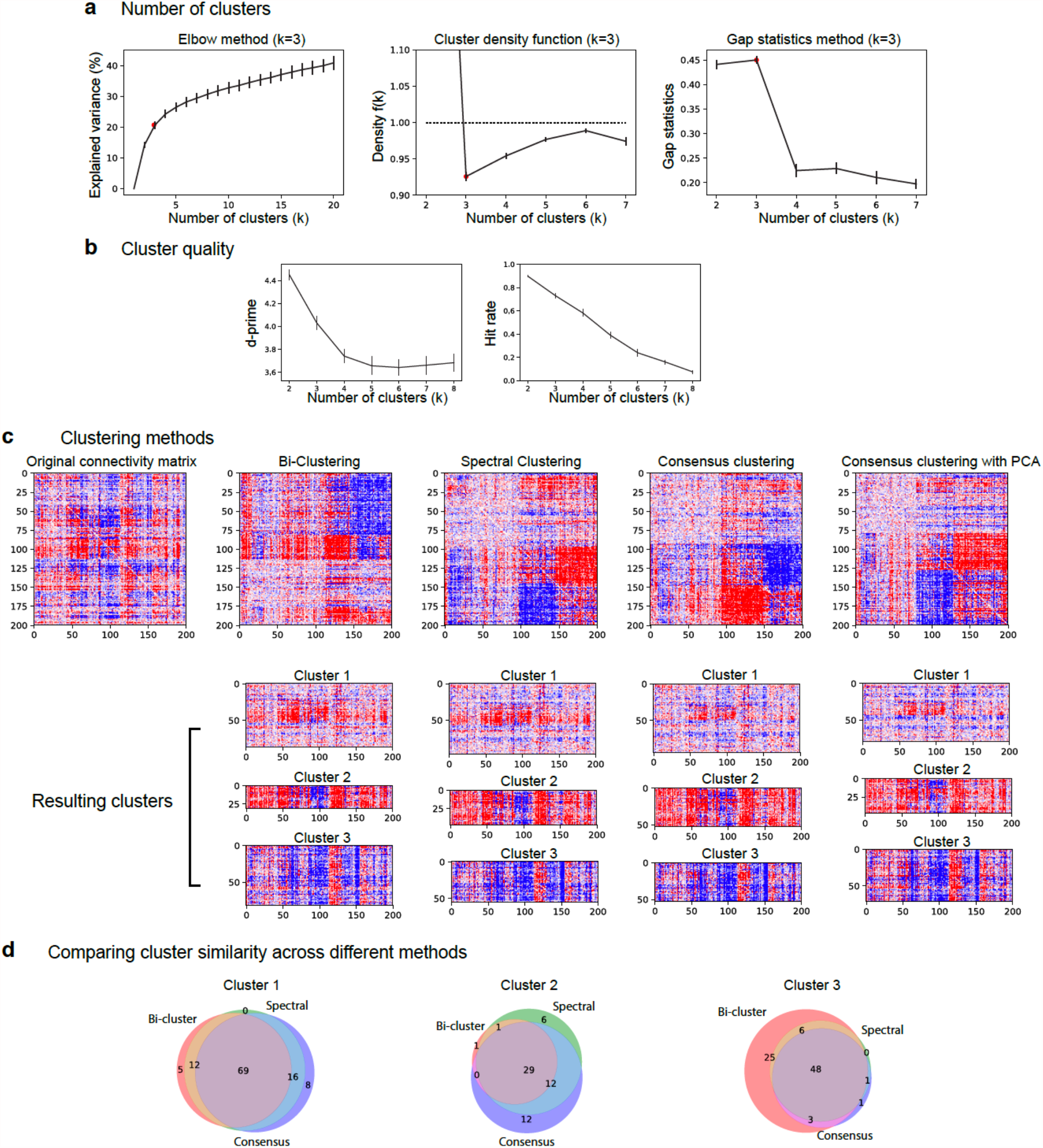
Number of clusters, cluster quality, and comparison of clustering methods. **a**) Number of clusters estimated with three different methods (Elbow method; cluster density function; Gap statistics; see Methods for details). Errorbars indicate standard error across mice (n=19). **b**) Quantification of cluster quality as the number of clusters and estimation kernels. The d-prime (d’) was calculated using Fisher’s linear discriminant analysis to find the line of maximum separation in PC space. The higher the value, the more distinguishable the clusters are. Hit-rate was calculated nearest-neighbors method (n_neighbors=3), which is a non-parametric estimate of exemplar contamination in each cluster. For each unit belonging to the cluster of interest, the three nearest units in principal-component space are identified. The “hit rate” is defined as the fraction of these units that belong to the cluster of interest. This metric is based on the “isolation” metric from (Hill et al., 2011). The higher the value, the less contamination in each cluster. Error bars indicate standard error across mice (n=19). **c**) Original functional connectivity matrix from an example mouse and four different clustering methods: bi-clustering, spectral clustering, consensus clustering on original matrix, and consensus clustering after PCA (the method used in this paper). The lower plots show the resulting clusters from different methods. **d**) Venn diagram illustrating the overlapping degree of different clusters estimated with different methods from the example mouse in c. This result suggests that our clustering findings are fundamental and the clustered units are roughly similar across clustering methods.

**Supplemental Figure 4.**
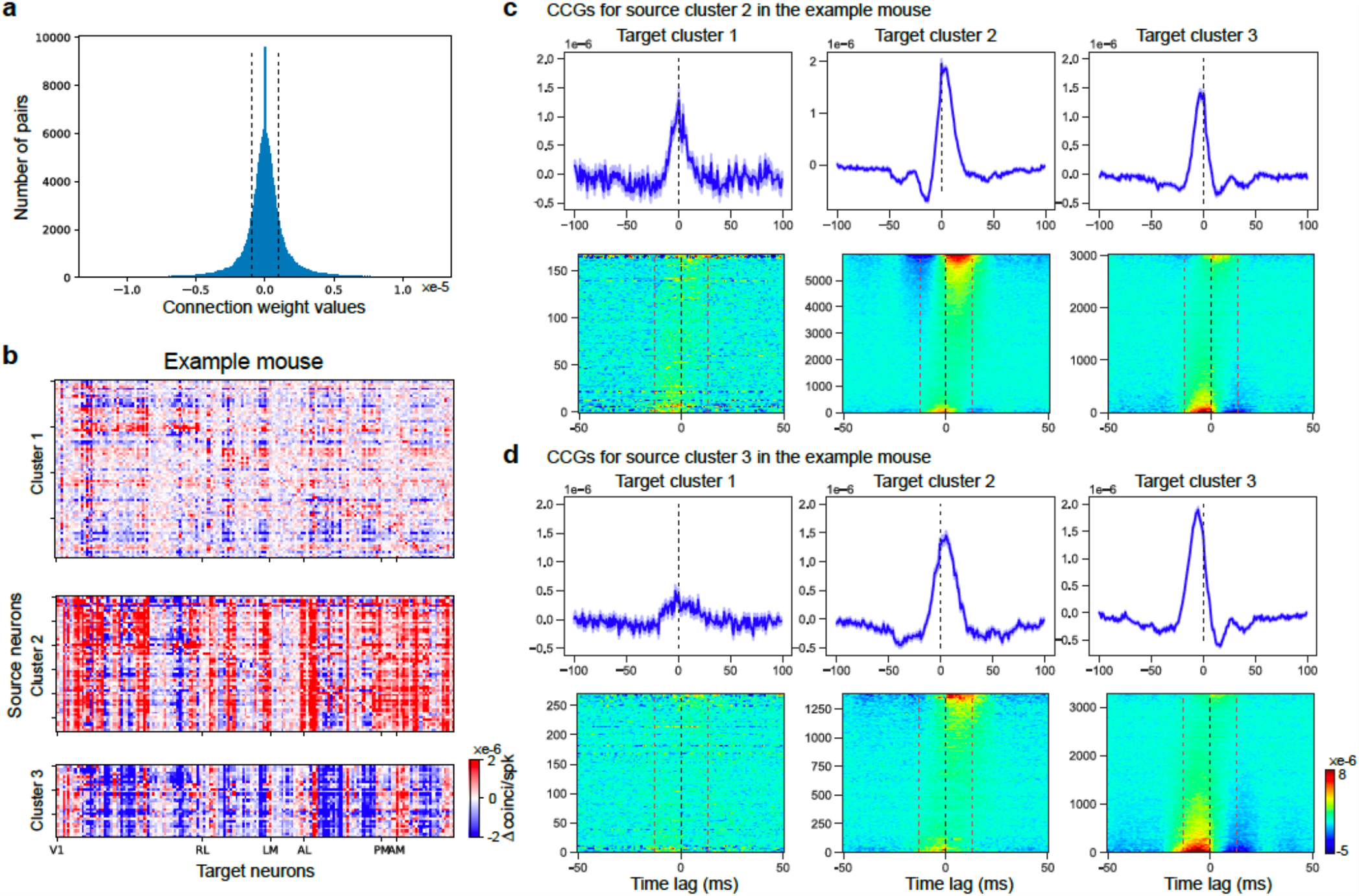
Shape of jitter-corrected CCGs for different sub-clusters. **a**) Distribution of connection weights of the connectivity matrix from all mice (n = 620,848 from 19 mice). Standard deviation of the distribution is 2.26*10^−6^ coincidence/spike. Dashed line indicates 1 std in the distribution. b) Clustering result from the example mouse in Fig. 1g. Cluster 1 is defined as source neurons with weak connections to the network. Cluster 2 is defined as source neurons with strong positive weight to the network. Cluster 3 is defined as source neurons with strong negative weight. Panels c) and d) illustrate the shape of jitter-corrected CCGs from the same example mouse in b). c) Jitter-corrected CCGs from source cluster 2 neurons to cluster 1 (n = 168 pairs), cluster 2 (n=5992 pairs) and cluster 3 (n=3024 pairs). Top: averaged CCGs for different groups. Bottom: waterfall plots of CCGs sorted by the calculated weight in the ±13 ms window of each CCG (highlighted by red dashed line) for different groups. Black dashed lines indicate 0 time lag. d) Jitter-corrected CCGs from source cluster 3 neurons to cluster 1 (n=270 pairs), cluster 2 (n=1380) and cluster 3 (n=3270). Errorbars indicate s.e.m.

**Supplemental Figure 5.**
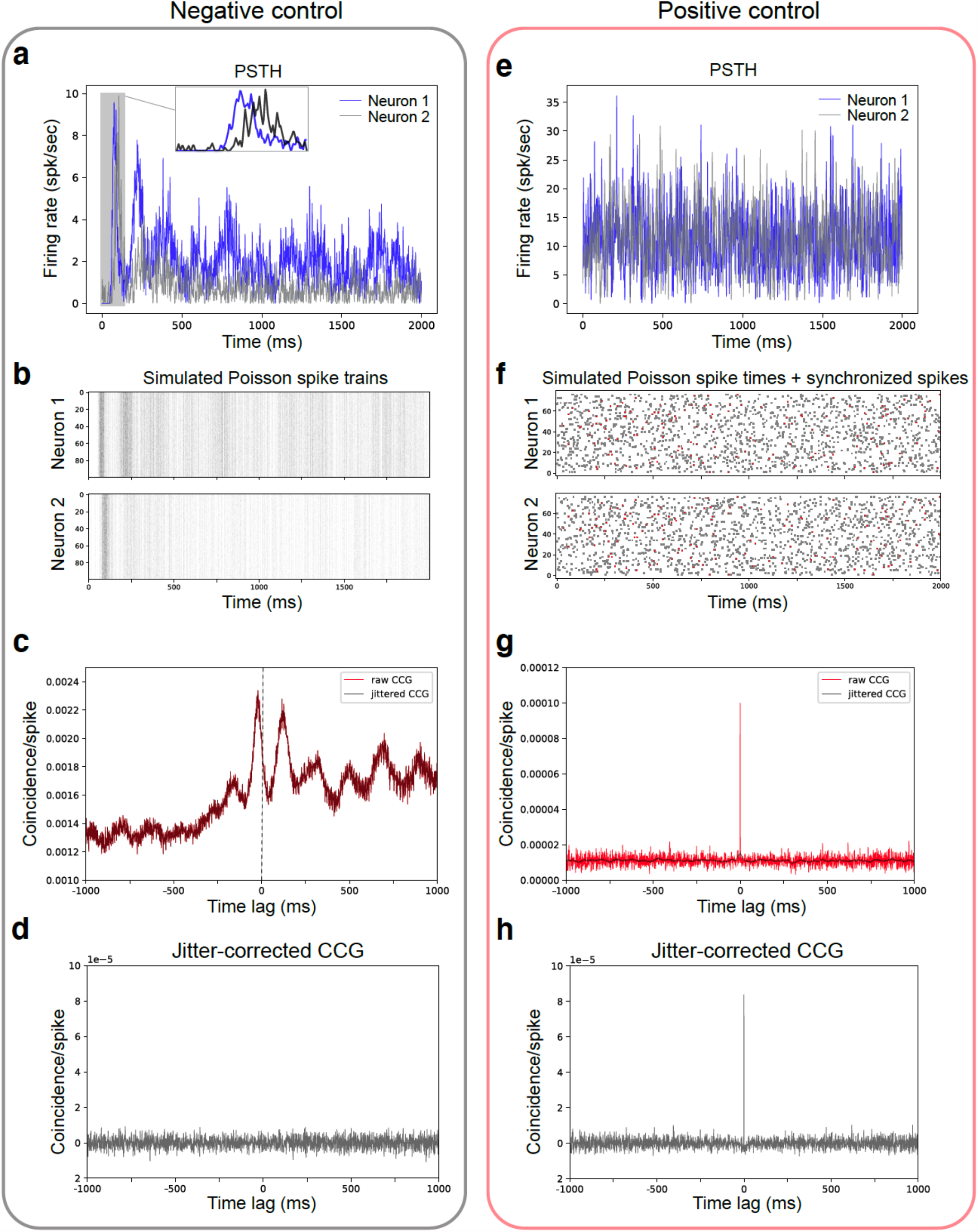
Testing the relationship between PSTH shape and CCG sharp peak. Two simulations were performed to test the relationship between the PSTH temporal dynamics of two neurons and their fast-timescale correlations, as reflected in the jitter-corrected CCG. The negative control was done by generating Poisson spike times with correlated PSTH temporal profiles; then, we tested whether there was a sharp peak in the jitter-corrected CCG with jitter window=25ms. **a**) PSTHs of two neurons (1 ms bin) with temporal Pearson’s correlation equal to 0.35. Neuron 1 has an onset earlier than neuron 2. **b**) Simulated Poisson spike trains based on the PSTHs in a) for the two neurons across repeats. **c**) Raw CCG (red line) and jittered CCG (calculated by jittering spike times large than 25 ms; black line) between neuron 1 and 2 (the probability of neuron 1 generating a spike relative to each spike in neuron 2) calculated based on the simulated Poisson spike times. Both CCGs showed a clear peak at a negative time lag, reflecting the fact that over long timescales, neuron 1 fires earlier than neuron 2, consistent with the temporal profile of the PSTHs. **d**) Jitter-corrected CCG calculated by subtracting jittered CCG from raw CCG in c). Because there is no fast timescale correlation within 25ms, predetermined by Poisson process, we didn’t observe any sharp peak in jitter-corrected CCG. This negative control indicates that a temporal correlation in the PSTH profile does not necessarily lead to a sharp peak in the CCG. The positive control was simulated by generating Poisson spike times given uncorrelated PSTH temporal profiles but adding additional synchronized spikes to the two simulated spike trains. Then, we tested whether the two spike trains showed any brief timescale correlation with jitter-corrected CCG. **e**) Uncorrelated PSTHs (Pearson’s r=0.01) of two neurons. **f**) Simulated Poisson spike train based on the PSTHs in e) with synchronized spikes (10% of total spikes; highlighted in red) added to both neuron 1 and 2. **g**) Raw and jittered CCGs between simulated spike trains. **h**) Jitter-corrected CCG calculated by subtracting jittered CCG from raw CCG. The jitter-corrected CCG showed a sharp peak at 0 time lag, indicating synchronization of spike times between the two neurons. This result is consistent with the injection of synchronized spikes, despite the fact that their PSTH profiles are uncorrelated.

